# Non-random sister chromatid segregation mediates rDNA copy number maintenance in *Drosophila*

**DOI:** 10.1101/498352

**Authors:** George J. Watase, Yukiko M. Yamashita

## Abstract

Although considered to be exact copies of each other, sister chromatids can segregate non-randomly in some cases. For example, sister chromatids of the X and Y chromosomes segregate non-randomly during asymmetric division of male germline stem cells (GSCs) in *D. melanogaster*. Here we identify that the ribosomal DNA (rDNA) loci, which are located on the X and Y chromosomes, and an rDNA-binding protein, Indra, are required for non-random sister chromatid segregation (NRSS). We provide the evidence that NRSS is a mechanism by which GSCs recover rDNA copy number, which occurs through unequal sister chromatid exchange, counteracting the spontaneous copy number loss that occurs during aging. Our study reveals an unexpected role for NRSS in maintaining germline immortality through maintenance of a vulnerable genomic element, rDNA.

**One Sentence Summary:** rDNA copy number maintenance by non-random sister chromatid segregation contributes to germline immortality in *Drosophila*

---

Sister chromatids, generated through the precise process of DNA replication, are considered identical. Nevertheless, it has been proposed that sister chromatids might carry distinct information or mutation loads, and their non-random segregation may underlie asymmetric cell division (*1-3*). However, the underlying mechanism remains elusive, preventing investigation of the physiological relevance of non-random sister chromatid segregation (NRSS). Using *D. melanogaster* male germline stem cells (GSCs) as a model system, where asymmetric stem cell division can be observed at single cell resolution, we previously showed that the X and Y chromosomes exhibit strikingly biased sister chromatid segregation (*4*). By using chromosome-orientation fluorescence *in situ* hybridization (CO-FISH) with chromosome-specific probes (Fig. 1A), (+)-strand templated vs. (-)-strand templated sister chromatids of each chromosome can be differentiated (Fig. 1A and B). If sister chromatids are equivalent, (+)-vs. (-)-strand templated sister chromatids would segregate to the GSC or GB (gonialblast, the differentiating daughter of a GSC) at random (50:50). Although we observed random sister chromatid segregation for autosomes (chromosome 2 and 3), the X and Y chromosomes segregated their sister chromatids non-randomly, with a specific strand segregating to GSC in ∼80% of observed divisions (Fig. 1C, ‘red strand’)(*4*). This demonstrated that sister chromatids, which supposedly carry the same genetic information, can be distinguished and segregated non-randomly during asymmetric stem cell division.

**Fig. 1.**
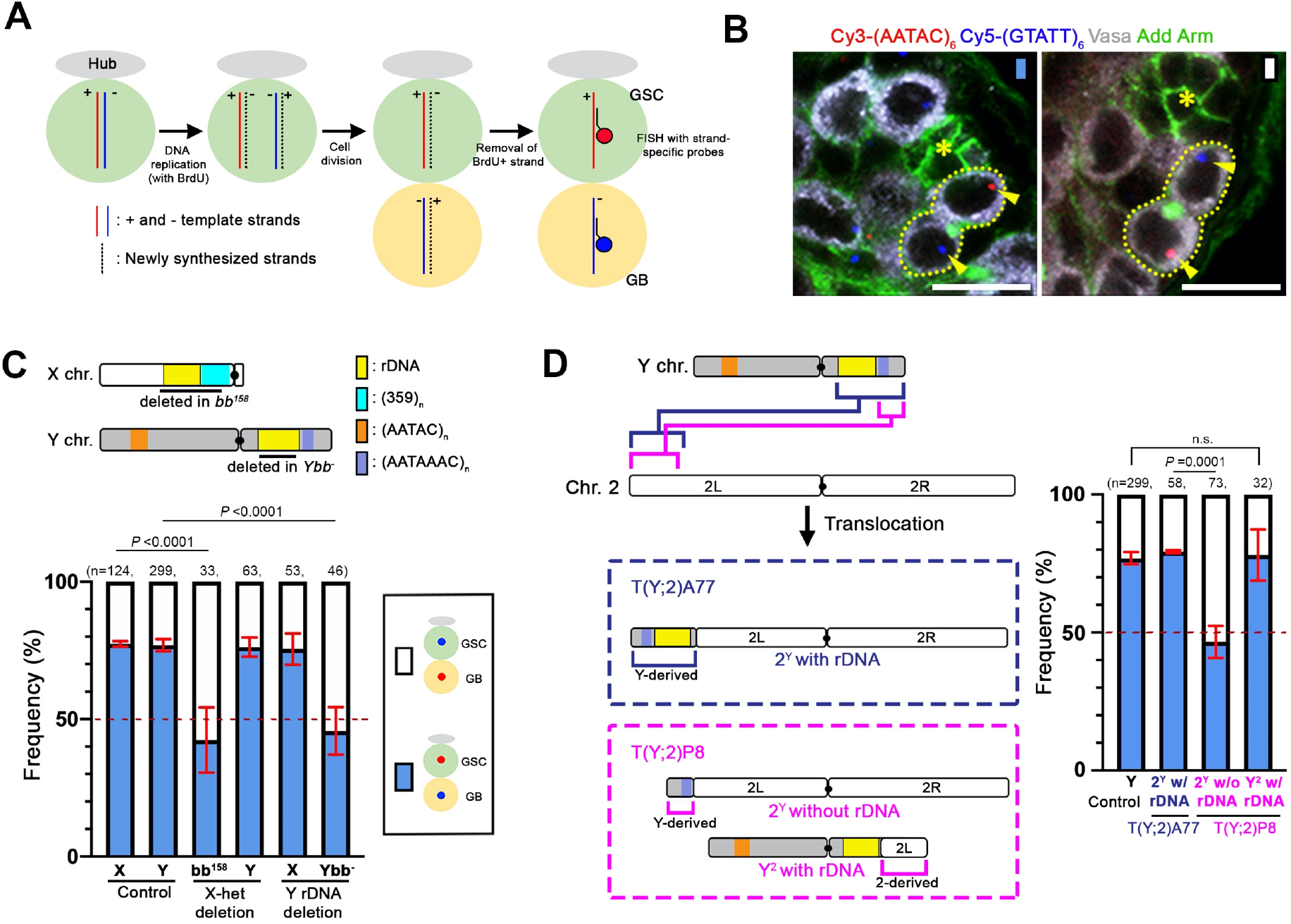
rDNA loci are required for NRSS of the X and Y chromosomes in *D. melanogaster*. (A) Chromosome-orientation *in situ* hybridization (CO-FISH) to assess NRSS. Plus (+) vs. minus (-) templated strands are indicated by red and blue lines, newly-synthesized strands by black dotted lines. Following removal of BrdU-containing, newly-synthesized strands, strand-specific probes were applied to distinguish red vs. blue templated strands. (B) Representative images of Y chromosome CO-FISH results where a GSC inherits the ‘red’ strand ((AATAC)_n_), whereas a GB inherits the ‘blue’ strand ((GTATT)_n_,. The hub, the stem cell niche to which GSCs are attached, is indicated by an asterisk, GSC-GB pairs are outlined by dotted lines and the CO-FISH signals by arrowheads. Vasa: germ cells. Arm: hub. Add: the connection between GSC and GB. Bar: 10µm. (C, D) Schematics of *D. melanogaster* X and Y chromosomes (C) and Y-2 translocation chromosomes (D). Summary of sister chromatid segregation patterns in indicated genotypes is shown (see table S1 and S2). Data shown as mean ± s.d. from three independent experiments. n, number of GSC-GB pairs scored. *P*-values: Fisher’s exact test.

To elucidate the underlying molecular mechanism of NRSS, we sought to identify the chromosomal loci that mediate NRSS and found that ribosomal DNA (rDNA) is required. An X chromosome without rDNA, *Df(1)bb*^*158*^ (*bb*^*158*^ hereafter), as well as a Y chromosome without rDNA (*Ybb*^*-*^), exhibited randomized sister chromatid segregation (Fig. 1C, table S1). Importantly, the intact Y chromosome in the *bb*^*158*^ strain, as well as the intact X chromosome in the *Ybb*^*-*^ strain, still exhibited NRSS (Fig. 1C, table S1), suggesting that the rDNA loci likely act as cis-elements to mediate NRSS. A chromosome 2 containing an rDNA locus translocated from the Y chromosome also exhibited NRSS (‘2^Y^ with rDNA’ in T(Y;2)A77 translocation, Fig. 1D, table S2), suggesting that rDNA is sufficient to induce NRSS. As a critical control, a chromosome 2 carrying a similar translocation from the Y chromosome that does not include the rDNA did not exhibit NRSS (‘2^Y^ without rDNA’ in T(Y;2)P8 translocation, Fig. 1D, table S2). This is the first demonstration that a specific region of a chromosome is responsible for NRSS, opposing the widely-held speculation that NRSS depends on chromosome-wide information such as epigenetic information and replication-induced mutations (*5*).

To understand how rDNA mediates NRSS, we isolated rDNA binding proteins from the GSC extract. Each rDNA locus consists of 150-225 repeated rDNA units in order to support the high demand of ribosome biogenesis (*6*). Each rDNA unit contains the 18S, 5.8S/2S, 28S rRNA genes and three spacer sequences [the external transcribed spacer (ETS), internal transcribed spacer (ITS) and intergenic spacer (IGS)] (Fig. 2A). Interestingly, the Y chromosome of *D. simulans*, a species closely related to *D. melanogaster*, has IGS repeats but no rRNA genes, ETS or ITS (*7*), yet exhibited NRSS (fig. S1, table S1). We hypothesized that IGS may be responsible for NRSS. Thus, we isolated IGS-binding proteins by mass spectrometry followed by a secondary screen based on subcellular localization (Fig. 2B, table S3). In this study, we focus on a previously-uncharacterized zinc finger protein, CG2199, which we named Indra after the Hindu god who lost immortality due to a curse from Durvasa. Using a specific anti-Indra antibody (fig. S2A) and an Indra-GFP line, we found that Indra localizes to the nucleolus (the site of rDNA transcription) in interphase (Fig. 2C, fig. S3A) and rDNA loci during metaphase (Fig. 2D, fig. S3B). ChIP-qPCR further demonstrated that Indra preferentially binds to IGS (Fig. 2E). Strikingly, RNAi-mediated knockdown of *indra* in the germline (fig. S2A) compromised NRSS for both the X and Y chromosomes (Fig. 2F, table S4). Taken together, these results show that IGS and its binding protein Indra mediate NRSS.

**Fig. 2.**
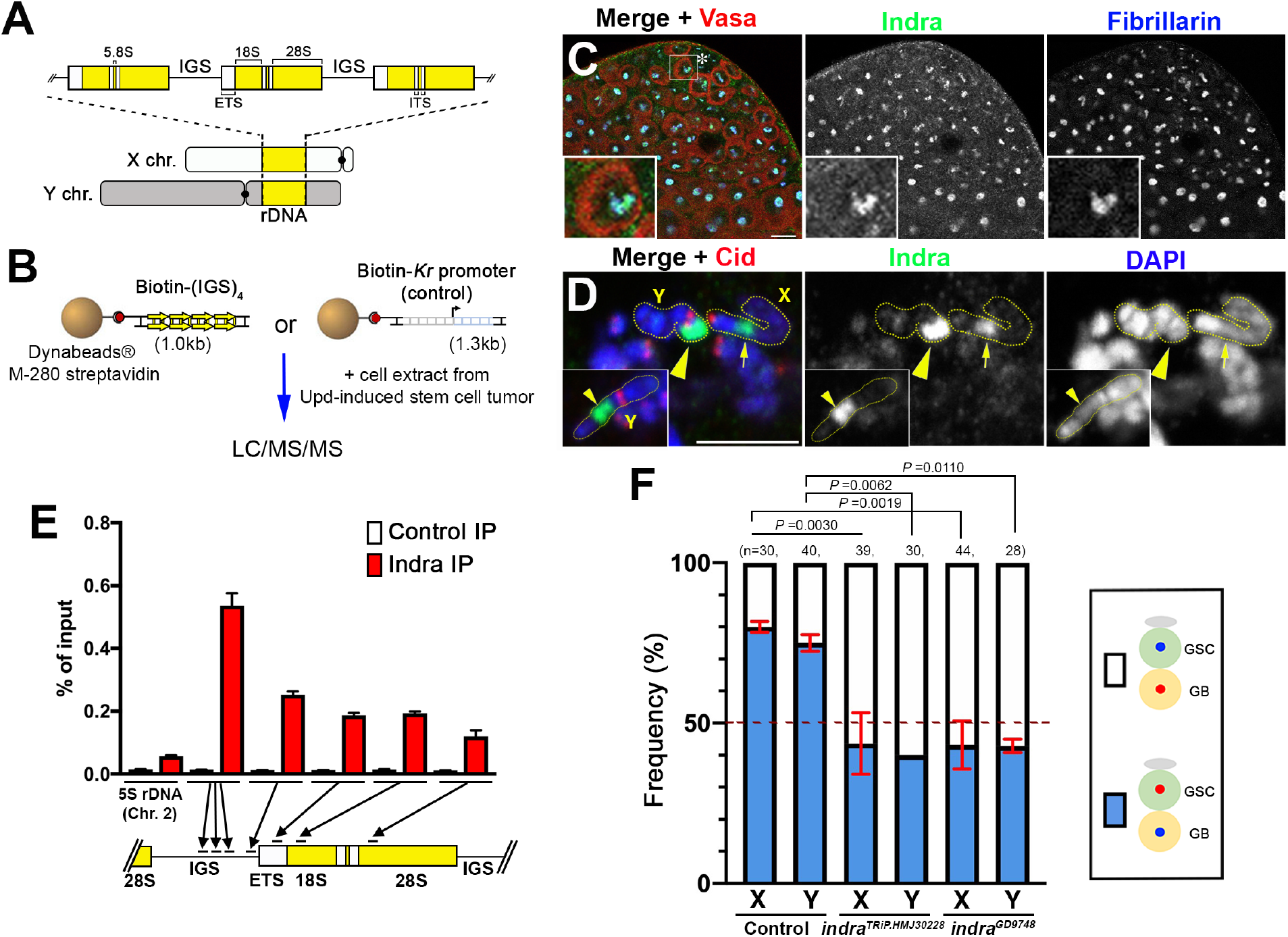
Indra is a novel zinc-finger protein that binds to rDNA and mediates NRSS. (A) Schematic of rDNA loci in *D. melanogaster*. (B) Experimental scheme to isolate IGS-binding proteins (see Materials and Methods). (C) Localization of Indra at the apical tip of the testis. The hub is indicated by an asterisk. An enlarged image of a GSC is shown in the inset. Fibrillarin: nucleolus. Vasa: germ cells. Bar: 10µm. (D) Localization of Indra on a metaphase chromosome spread from germ cells. The X rDNA locus (arrow) and Y rDNA locus (arrowhead) can be identified by their relative location to the centromere (Cid). An additional example of the Y chromosome is shown in the inset. Bar: 5 µm. (E) Indra ChIP-qPCR showing enrichment of Indra on rDNA/IGS. The 5S rDNA sequence on chromosome 2 outside of the rDNA loci was used as a negative control. Mean and s.d. from three technical replicates of qPCR are shown. Similar results were obtained from two biological replicates. (F) Summary of sister chromatid segregation patterns upon knockdown of *indra* (see table S4). Data shown as mean ± s.d. from three independent experiments. n, number of GSC-GB pairs scored. *P*-values: Fisher’s exact test.

We found that *indra* is required for rDNA copy number maintenance. RNAi-mediated depletion of *indra* (*nos-gal4>UAS-indra*^*TRiP*.*HMJ30228*^) resulted in drastically fewer progeny compared to control (Fig. 3A, P_0_). Some of the offspring from *indra*^*TriP*.*HMJ30228*^ males exhibited a *bobbed* phenotype, a hallmark of rDNA copy number insufficiency characterized by abnormal cuticle patterns on the abdomen (Fig. 3B; (*8*)). The frequency of *bobbed* flies increased when the Y chromosome from *indra*^*TriP*.*HMJ30228*^ fathers was placed in the background of *bb*^*158*^, the X chromosome that lacks rDNA (Fig. 3B). Quantitative droplet digital PCR (ddPCR) confirmed that rDNA copy number was reduced in *indra*^*TriP*.*HMJ30228*^ animals (Fig. 3C, P_0_). Depletion of *indra* over successive generations resulted in a progressive loss of fecundity (Fig. 3A, F_1_-F_2_) associated with a reduction in rDNA copy number (Fig. 3C, F_1_-F_2_). Moreover, *indra* is required for ‘rDNA magnification’, a phenomenon by which an X chromosome with insufficient rDNA copy number is induced to recover copy number, when the fly lacks rDNA on the Y chromosome (*Ybb*^*-*^) (Fig. 3, D and E, fig. S4; (*9*)). These results suggest that *indra* is required for rDNA copy number maintenance over generations.

**Fig. 3.**
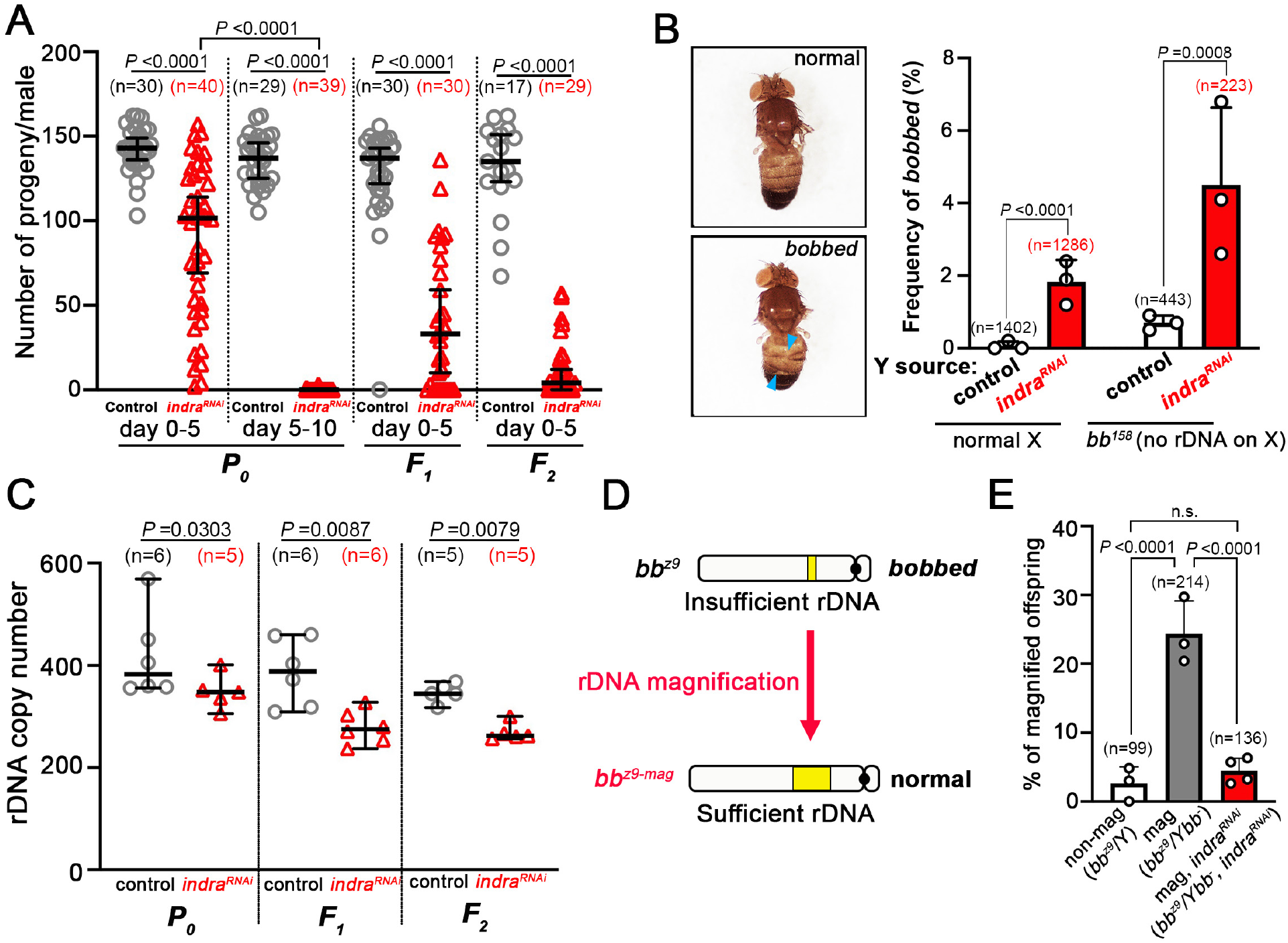
*indra* is required for rDNA copy number maintenance. (A) Fertility of control and *nos-gal4*>*UAS-indra*^*TRiP*.*HMJ30228*^ males across generations (P_0_-F_2_). Total number of progeny from 0-5 day old males in each generation and 5-10 day old males in P_0_ were scored. Data shown as median with 95% confidence interval and individual data points. n, number of individual crosses scored. *P*-value: two-tailed Mann-Whitney test. (B) Frequency of *bobbed* animals in progeny of 0-5 day-old control and *nos-gal4>UAS-indra*^*TRiP*.*HMJ30228*^ males. Mean and s.d. from three independent experiments with individual data points are shown. n, total number of progeny scored. *P*-values: two-tailed chi-squared test. Examples of normal and ‘*bobbed*’ cuticle phenotypes are shown on the left. (C) 28S rRNA gene copy number in the testes from 0-5 day old control and *nos-gal4*>*UAS-indra*^*TRiP*.*HMJ30228*^ males in successive generations (P_0_-F_2_) assessed by ddPCR. Data shown as median with 95% confidence interval and individual data points. n, number of individual crosses scored. *P*-value: two-tailed Mann-Whitney test. (D) Schematic diagram of rDNA magnification assay (see fig. S4 for the details). Magnification was detected by normal cuticle phenotype in the offspring. (E) Frequency of rDNA magnification in the indicated genotypes/conditions. Data shown as mean and s.d. from four (*nos-gal4*>*UAS-indra*^*GD9748*^, *UAS-Dcr-2*) or three (the rest) independent experiments with individual data points. n, total number of progeny scored. *P*-values: Fisher’s exact test.

Although the repetitiveness of rDNA loci is critical to support ribosome biogenesis, it also makes rDNA loci susceptible to intrachromatid recombination, which leads to spontaneous copy number loss (Fig. 4A). To maintain the integrity of rDNA loci, copy number loss must be counteracted by copy number recovery. In yeast, rDNA copy number recovery is mediated by unequal sister chromatid recombination (*10*). Similarly, rDNA magnification, which we postulated to mediate rDNA copy number recovery in the *Drosophila* male germline (*11*), is proposed to utilize unequal sister chromatid exchange (USCE) (*9*). USCE allows for copy number recovery on one of the sister chromatids at the expense of the other (Fig. 4B; (*9*)), generating asymmetry between two sister chromatids (Fig. 4B). We hypothesized that this asymmetry may underlie NRSS.

**Fig. 4.**
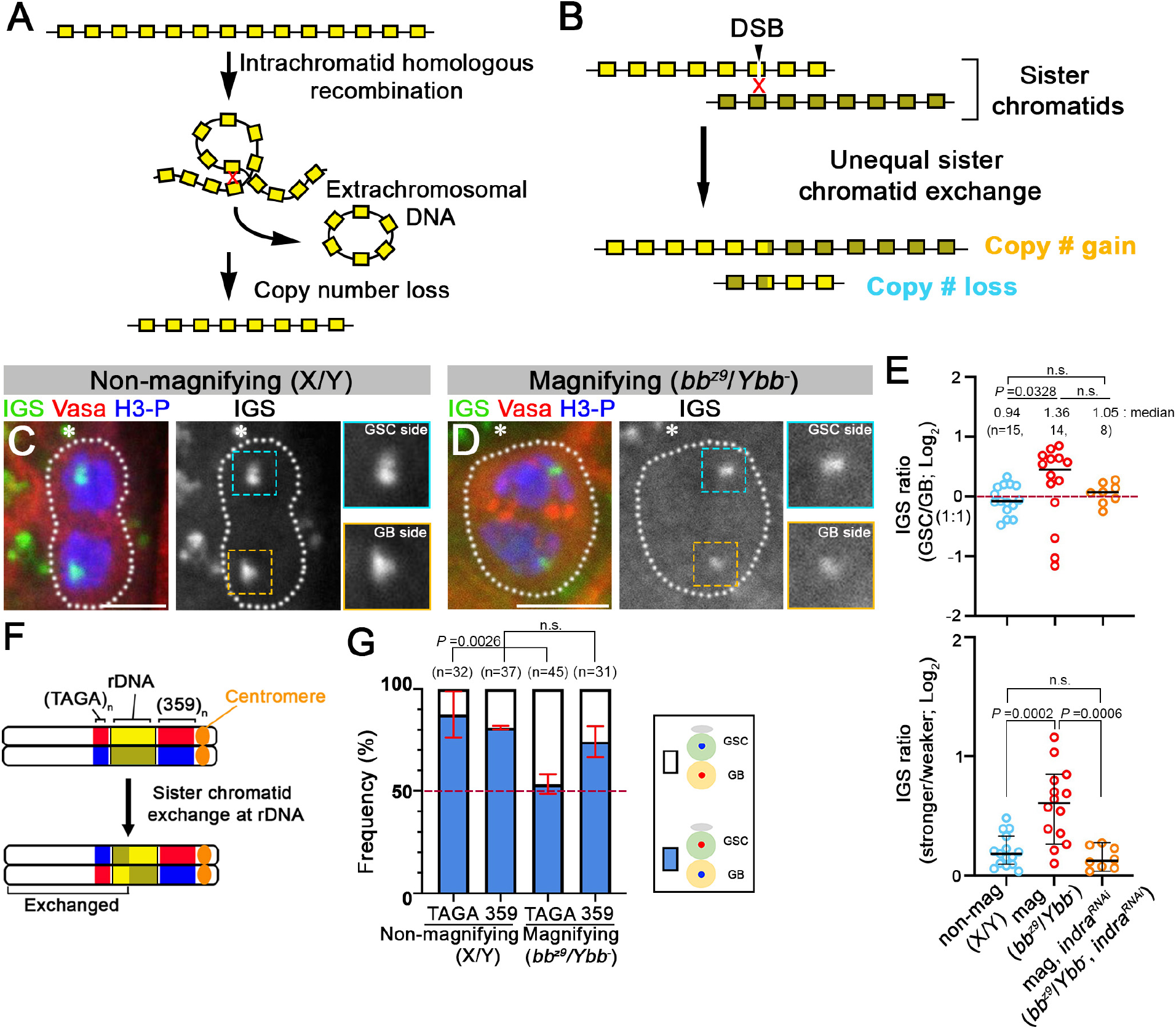
rDNA loci undergoe sister chromatid exchange and segregate asymmetrically in GSCs during rDNA magnification. (A) Diagram of spontaneous rDNA copy number loss by intrachromatid recombination. (B) A proposed model for rDNA copy number recovery by unequal sister chromatid exchange. (C, D) DNA-FISH for IGS during anaphase in GSCs under normal (*yw*) (C) and magnifying (*bb*^*z9*^*/Ybb*^*-*^) (D) conditions. The hub is indicated by an asterisk. An enlarged image of the IGS signal from GSC and GB sides is shown in the inset. Bar: 5 µm. (E) Quantification of IGS signal intensity presented as GSC side/GB side (top panel) or stronger/weaker (bottom panel) during anaphase in GSCs in control and *nos-gal4*>*UAS-indra*^*GD9748*^, *UAS-Dcr-2* males. The median and individual data point are shown. 95% confidence interval is also shown in the bottom panel. n, number of anaphase cells scored. *P*-value: two-tailed Mann-Whitney test. Note that due to rare cases where the GB side exhibited stronger IGS signal than GSC side, the data did not reach statistical significance between control and *indra*^*GD9748*^ under magnifying conditions in the top panel. (F) Diagram of the X chromosome showing the location of the rDNA and the 359-bp and (TAGA)_n_ repeats. Sister chromatid exchange at rDNA loci would flip (TAGA)_n_ segregation pattern relative to 359-bp. (G) Summary of sister chromatid segregation patterns assessed by 359-bp and (TAGA)_n_ probes in the indicated genotypes/conditions (see table S5). Data shown as mean ± s.d. from three independent experiments. n, number of GSC-GB pairs scored. *P*-values: Fisher’s exact test.

Strikingly, asymmetry in rDNA amount was detected during anaphase in GSCs under ‘magnifying conditions’(*bb*^*z9*^*/Ybb*^*-*^) with the GSCs preferentially inheriting the stronger signal (Fig. 4, D and E). As an important control, asymmetry in rDNA amount was not observed in flies with sufficient rDNA copy number (Fig. 4, C and E). Interestingly, there were rare cases where rDNA asymmetry was created, but the stronger signal was inherited by the GBs (Fig. 4E, top panel). Plotting the ratio of stronger over weaker rDNA signal (Fig. 4E, bottom panel) suggests that the magnifying condition strongly induced USCE. rDNA copy number and segregation asymmetries were absent under magnifying conditions following depletion of *indra* (Fig. 4E), suggesting that *indra* may be involved not only in NRSS but also in generating copy number asymmetry through USCE.

These results are consistent with a model in which rDNA magnification is mediated by USCE, and the sister chromatid with increased rDNA copy number is selectively retained by GSCs by NRSS. We further tested this idea by using additional CO-FISH probes. The probe for the 359-bp repeat, located proximal to the rDNA (Fig. 4F), was used to detect NRSS in the experiments described above. (TAGA)_n_, which is located distal to the rDNA, also exhibited NRSS in control (non-magnifying) conditions (Fig. 4G, table S5). Strikingly, under magnifying conditions, 359-bp maintained NRSS but (TAGA)_n_ exhibited random segregation, suggesting that sister chromatid exchange occurred between 359-bp and (TAGA)_n_, most likely within the rDNA locus (Fig. 4F). Taken together, these data suggest that GSCs undergo rDNA magnification through USCE followed by NRSS (see Supplementary text).

Our study reveals the unexpected molecular mechanisms and biological significance of NRSS. We propose that NRSS is a key process to recover and maintain inherently unstable rDNA copy numbers such that the integrity of the germline genome is upheld over generations, supporting germline immortality. Future work is required to understand how rDNA copy number differences between sister chromatids are recognized and faithfully segregated to the GSCs to achieve rDNA copy number recovery.

## Competing interests

The authors declare no competing interests.

## Acknowledgements

We thank the Bloomington, Kyoto, and National Drosophila Species Stock Centers, the Vienna Drosophila Resource Center and the Developmental Studies Hybridoma Bank for reagents. We thank the Yamashita lab members, Drs. Swathi Yadlapalli and Sue Hammoud, and the Life Science Editors for comments on the manuscript. We thank MS Bioworks for mass-spectrometry analysis and the laboratories of Drs. Stephen Weiss and Jiandie Lin for sharing equipment. This research was supported by Howard Hughes Medical Institute.

## Author contributions

G.W. and Y. Y. designed and conducted experiments, interpreted results and wrote and edited the manuscript.

## Supplementary Materials

Materials and Methods

Supplementary Text

Figs. S1 to S5

Tables S1 to S7

## Materials and method

### Fly husbandry and strains

All fly stocks were raised on standard Bloomington medium at 25°C containing 0.15% of tegosept as anti-fungal (no propionic acid was added). The following fly stocks were used: *Df(1)bb*^*158*^, *y*^*1*^*/Dp(1;Y)y*^*+*^*/C(1)*; ca*^*1*^ *awd*^*K*^ (BDSC3143), *FM6/C(1)DX, y* f*^*1*^*/Y* (BDSC784), *UAS-indra*^*TRiP*.*HMJ30228*^ (BDSC63661), *UAS-Dcr-2* (BDSC24650), *indra-GFP* (BDSC67660; http://flybase.org/reports/FBti0186577) were obtained from the Bloomington Drosophila Stock Center. *y*^*1*^ *eq*^*1*^*/Df(YS)bb*^*-*^ (DGRC101260), *T(Y;2)A77, B*^*S*^, *y*^*+*^*/SM1; C(1)RM, y*^*1*^*/C(1;Y)1, y*^*1*^ (DGRC130079), *T(Y;2)P8, B*^*S*^, *y*^*+*^*/SM1; C(1)RM, y*^*1*^*/C(1;Y)1, y*^*1*^ (DGRC130170) were obtained from the Kyoto Stock Center. *D. simulans W*^*501*^(DSSC14021-0251.195) was obtained from the National Drosophila Species Stock Center. *UAS-indra*^*GD9748*^ (v20839) was obtained from the Vienna Drosophila Resource Center. *nos-gal4* (*1*), *UAS-Upd* (*2*), *tub-gal80*^*ts*^ (*3*), *nos-gal4* without VP16 (*4*) have been previously described.

To examine the sister chromatid segregation patterns of the 2^Y^ and Y^2^ chromosomes, *T(Y;2)A77/SM1; C(1)RM/O* or *T(Y;2)P8/SM1; C(1)RM/O* females were crossed to *yw* males and the resulting *T(Y;2)A77/+; X/O* and *T(Y;2)P8/+; X/O* male flies were examined. The details of the translocation are shown in Fig. 1D.

Two independent RNAi lines, *UAS-indra*^*TRiP*.*HMJ30228*^ and *UAS-indra*^*GD9748*^, were used to knockdown *indra* specifically in early germ cells using *nos-gal4* as the driver. *UAS-indra*^*GD9748*^ was combined with *UAS-Dcr-2* to increase RNAi efficiency. The knockdown efficiency of these RNAi lines was validated by immunostaining using an anti-Indra antibody (fig. S2A). Since *nos-gal4>UAS-indra*^*TRiP*.*HMJ30228*^ results in severe germ cell loss due to high RNAi efficiency (Fig. 3A, fig. S2A), a temperature-sensitive GAL4 inhibition system (*tub-gal80*^*ts*^; *nos-gal4τ1VP16>UAS-indra*^*TRiP*.*HMJ30228*^) was used as necessary (e.g. Fig. 2F). Upon shifting from the permissive temperature (18°C) to the non-permissive temperature (29°C), GSCs were lost gradually over 2-4 days (fig. S2, B and C), and the CO-FISH assay (Fig. 2F) was conducted 3 days after temperature shift. In assays that required a sustained germline (e.g. magnification assays, Fig. 3E and Fig. 4E), we used *nos-gal4>UAS-indra*^*GD9748*^, *UAS-Dcr-2*.

### Immunofluorescence staining and confocal microscopy

*Drosophila* adult testes were dissected in phosphate-buffered saline (PBS), transferred to 4% formaldehyde in PBS and fixed for 30 min. The testes were then washed in PBST (PBS containing 0.1% Triton X-100) for at least 30 min, followed by incubation with primary antibody in 3% bovine serum albumin (BSA) in PBST at 4°C overnight. Samples were washed for 60 min (3 × 20 min washes) in PBST, incubated with secondary antibody in 3% BSA in PBST at 4°C overnight, washed as above, and mounted in VECTASHIELD with 4’,6-diamidino-2-phenylindole (DAPI; Vector Labs, Burlingame, CA). To examine Indra localization on mitotic chromosome spreads, *Drosophila* 3rd instar larval testes were dissected in PBS, transferred to 0.5% sodium citrate and incubated for 10 min, fixed in 4% formaldehyde in PBS for 4 min, then squashed between the cover slip and slide glass. The sample was frozen in liquid nitrogen, the cover slip was removed, and immediately washed in PBS, followed by immunofluorescence staining as described above, except that the incubation was performed on the slide glass in a humid chamber with the sample covered with a small piece of parafilm.

The primary antibodies used were as follows: rabbit anti-Vasa (1:200; d-26; Santa Cruz Biotechnology, Santa Cruz, CA), mouse anti-Adducin-like [1:20; 1B1; developed by H.D. Lipshitz, obtained from Developmental Studies Hybridoma Bank (DSHB)](*5*), mouse anti-Armadillo (1:100; N2 7A1; developed by E. Wieschaus, obtained from DSHB)(*6*), rat-anti Vasa (1:20; developed by A.C. Spradling and D. Williams, obtained from DSHB), mouse anti-Fibrillarin (1:200; 38F3; Abcam), chicken anti-Cid (1:500)(*7*). The anti-Indra antibody was generated by injecting a peptide (RKITDVLETITHRSIPSSLPIKIC) into guinea pig (Covance, Denver, PA) and used at a dilution of 1:500. Specificity of the antibody was validated by the lack of signal in *indra*^*RNAi*^ testis (fig. S2A). Alexa Fluor-conjugated secondary antibodies (Life Technologies) were used at a dilution of 1:200. Images were taken on a Leica TCS SP8 confocal microscope with a 63x oil immersion objective (NA = 1.4) and processed using Adobe Photoshop software.

For DNA FISH combined with immunofluorescent staining, whole mount *Drosophila* testes were prepared as described above, and the immunofluorescence staining protocol was carried out first. Upon completion of the wash post incubation with the secondary antibody, samples were fixed with 4% formaldehyde for 10 min and washed in PBST for 30 min. Fixed samples were incubated with 2 mg/ml RNase A solution at 37°C for 10 min, then washed with PBST. Samples were washed in 2x SSC with increasing formamide concentrations (20% and 50%) for 10 min each. Hybridization buffer (50% formamide, 10% dextran sulfate, 2x SSC, 1 mM EDTA, 1 mM probe) was added to washed samples. Samples were denatured at 91°C for 2 min, then incubated overnight at 37°C. Following the hybridization, testes were washed once in 50% formamide/2x SSC, once in 20% formamide/2x SSC, and 3 times in 2x SSC. All reagents contained 1 mM EDTA except for the washes prior to RNase A treatment. Fluorescence quantification was done on merged z-stacks using Image J ‘Sum of pixel intensity (RawIntDen)’ to compare signal intensity between sister chromatids. To avoid the effect of signal intensity changes along the Z plane, we scored anaphase GSCs only when two IGS FISH signals were found within the same Z plane for Fig. 4, C-E.

### Chromosome orientation fluorescence *in situ* hybridization (CO-FISH)

CO-FISH in whole mount *Drosophila* testes was performed as previously described (*8*). Briefly, young adult flies (day 1-3) were fed with 5-bromodeoxyuridine (BrdU)-containing food (950 µl of 100% apple juice, 7 mg of agar, and 50 µl of 100 mg/ml BrdU solution in a 1:1 mixture of acetone and DMSO) for 12 hours. After the feeding period, flies were transferred to regular fly food for 13.5 hours. Because the average GSC cell cycle length is ∼12 hour, most GSCs undergo a single S phase in the presence of BrdU followed by mitosis during this feeding procedure. GSCs that have undergone more or less than one S phase or mitosis were excluded from our analysis by limiting the scoring to GSC-GB pairs that have complementary CO-FISH signals in the GSC and GB (red signal in one cell, blue signal in the other). Note that GSC and GB stay connected by the fusome until mid-S phase, which allowed identification of the GSC-GB sister pairs. Testes were dissected, fixed and immunostained as described above. Then, testes were fixed for 10 min with 4% formaldehyde in PBS, followed by 3 washes with PBST. Following the washes, the testes were rinsed once with PBST and treated with RNase A (Roche; 2 mg/ml in PBS) for 10 min at 37°C, washed with PBST for 5 min, and stained with 100 µl of 2 µg/ml Hoechst 33258 (Invitrogen) in 2x SSC for 15 min at room temperature. The testes were then rinsed 3 times with 2x SSC, transferred to a tray, and irradiated with ultraviolet light in a CL-1000 Ultraviolet Crosslinker (UVP; wavelength: 365 nm; calculated dose: 5400 J/m^2^). Nicked BrdU positive strands were digested with exonuclease III (New England BioLabs) at 3 U/µl in 1x NEB1 buffer or 1x NEB cutsmart buffer for 10 min at 37°C. The testes were washed once with PBST for 5 min and then fixed with 4% formaldehyde in PBS for 2 min. Subsequently, the fixed testes were washed 3 times with PBST. Testes were incubated sequentially for a minimum of 10 min each in 20% formamide/2x SSC and 50% formamide/2x SSC. The testes were incubated with hybridization buffer (50% formamide, 2x SSC, 10% dextran sulfate) containing 1 µM of each probe for 16 hours at 37°C. Following hybridization, testes were washed once in 50% formamide/2x SSC, once in 20% formamide/2x SSC and 3 times in 2x SSC. Images were taken on either a Leica TCS SP5 or STELLARIS 8 confocal microscope with a 63x oil immersion objective (NA = 1.4) and processed using Adobe Photoshop software. For CO-FISH in GSCs from *tub-gal80*^*ts*^; *nos>indra*^*TRiP*.*HMJ30228*^, the BrdU pulse was conducted 3 days after temperature shift (fig. S2C). BrdU was fed at 29°C for 9 hours, followed by an 11-hour chase at 29°C in which the flies were fed regular fly food. The probes are described in table S6. All reagents contained 1 mM EDTA except for the washes immediately preceding an enzymatic reaction (RNase A and exonuclease III).

### IGS DNA pull down and mass-spectrometry

200 pairs of *upd*-expressing testes (*nos-gal4>UAS-upd*) were dissected in Schneider’s *Drosophila* Medium (Gibco) and washed 3 times with ice-cold PBS. *upd* expression causes overproliferation of GSC-like cells. The testes were homogenized in lysis buffer [20 mM Tris-HCl pH8.0, 1 mM EDTA, 10% Glycerol, 0.2% NP-40, 1 mM DTT, 1x solution of PhoSTOP cocktail (Roche), 1x solution of c0mplete EDTA-free protease inhibitor cocktail (Roche)], and the homogenate was incubated on ice for 20 min. Following this incubation, the lysate was centrifuged at 3,000 rpm for 10 min at 4°C, and the supernatant was saved as whole cell extract. The pellet, which contains the nuclear fraction, was resuspended in lysis buffer containing 100 mM NaCl and incubated on ice for 1 hour. During incubation, the sample was vortexed at highest setting for 15 sec every 10 min. The nuclear fraction was isolated by centrifugation at 14,000 rpm for 30 min at 4°C and mixed with the whole cell extract prepared above. Protein concentration was measured by absorbance at 562 nm using Pierce™ BCA Protein Assay Kit (Thermo Scientific) to adjust the concentration between samples.

The 240-bp IGS sequence (repeated 4 times, 4xIGS), or the *Kr* gene promoter sequence (control), was cloned into pBluescript SK^-^. Biotin end-labeling at the 5’ of one strand of the 4xIGS or *Kr* gene promoter was performed by PCR using a T7 primer with Biotin-TEG and a T3 primer. Biotinylated 4xIGS and *Kr* gene promoter DNA were then purified by QIAGEN’s PCR purification kit. 2 µg of each biotinylated DNA was immobilized to 100 µl of streptavidin-bound M-280 Dynabeads™ (invirogen). The beads were washed 3 times with 1x Binding and Washing buffer (5 mM Tris-HCl pH8.0, 0.5 mM EDTA, 500 mM NaCl) and then blocked with 0.5% BSA in TGEDN buffer (120 mM Tris-HCl pH8.0, 1 mM EDTA, 100 mM NaCl, 1 mM DTT, 0.1% Triton X-100, 10% Glycerol). 20% volume of each of the biotinylated DNA-conjugated Dynabeads™ was incubated with 20 µg of herring sperm DNA (Sigma-Aldrich) and the cell extract prepared above (containing 500 µg of protein at 3.8-4.8 µg/µl concentration, matched between control vs. IGS beads). After incubating for 2 hours at 4°C, the beads were washed 5 times with TGEDN buffer. The proteins bound to either the 4xIGS or *Kr* gene promoter DNA were eluted in LDS sample loading buffer (1.5x) at 100°C for 15 min. 50% volume of each DNA bound proteins was separated on a 10% Bis-Tris Novex mini-gel (invitrogen) using the MES buffer system. The gel was stained with coomassie and excised into ten equally sized segments. These segments were analyzed by LC/MS/MS (MS Bioworks, Ann Arbor, MI). The gel digests were analyzed by nano LC/MS/MS with a Waters NanoAcquity HPLC system interfaced to a Thermo Fisher Q Exactive. Peptides were loaded on a trapping column and eluted over a 75 µm analytical column at 350 nL/min; both columns were packed with Luna C18 resin (phenomenex). The mass spectrometer was operated in data-dependent mode, with MS and MS/MS performed in the Orbitrap at 70,000 FWHM resolution and 17,500 FWHM resolution, respectively. The fifteen most abundant ions were selected for MS/MS.

### ChIP-qPCR

200 pairs of *upd*-expressing testes (*nos-gal4>UAS-upd*) were dissected in ice-cold PBS containing protease inhibitor [1x solution of c0mplete protease inhibitor cocktail (Roche) and 1 mM PMSF]. The testes were crosslinked by incubating with 1% formaldehyde for 15 min at 37°C and rinsed twice in ice-cold PBS containing protease inhibitor to stop the crosslink reaction. The testes were homogenized in 200 µl of ice-cold ChIP Sonication Buffer [1% triton X-100, 0.1% sodium deoxycholate, 50 mM Tris-HCl (pH 8.0), 150 mM NaCl, 5 mM EDTA], and the homogenate was incubated on ice for 15 min. Following the incubation, the homogenate was aliquoted into 0.5 ml PCR tubes, placed in a Biorupter^®^ Plus sonication system (DIAGENODE) and sonicated in 4°C water bath for 10 cycles of 30 sec ‘ON’ and 30 sec ‘OFF’ at ‘HIGH’ setting. The sonicated lysate was centrifuged at 14,000 rpm for 10 min at 4°C to pellet cell debris. The volume of supernatant was brought up to 1 ml with ChIP sonication buffer, and 40 µl of Dynabeads™ Protein A (invitrogen) was added to the supernatant. After a 1-hour preabsorption with Dynabeads™ Protein A at 4°C, 30 µl of supernatant (3%) was kept as ‘input’. The rest was split into two and incubated overnight with 10 µl of anti-Indra antibody (1:10 dilution from the original serum; generated as described above) or 10 µl of pre-immune guinea pig serum (1:10 dilution from the original serum), respectively. After incubating for 16 hours, 40 µl Dynabeads™ Protein A was added to each reaction and incubated for an additional 4 hours at 4°C with rotation. The beads were then washed for 5 min at 4°C with 1 ml of the following buffers: 2 washes with ChIP sonication buffer, followed by 3 washes with High Salt Wash buffer [1% triton X-100, 0.1% sodium deoxycholate, 50 mM Tris-HCl (pH 8.0), 500 mM NaCl, 5 mM EDTA]; 2 washes with LiCl Immune Complex Wash buffer [250 mM LiCl, 0.5% NP-40, 0.5% deoxycholate, 1 mM EDTA, 10 mM Tris-HCl (pH8.0)]; 1 wash with TE buffer [10 mM Tris-HCl (pH8.0), 1 mM EDTA). For elution, each ChIP sample was incubated with 250 µl of Elution Buffer (1% SDS, 100 mM NaHCO_3_) for 30 min at 65°C, vortexing gently every 10 min. After repeating the elution process once more, the supernatants were combined. 500 µl of elution buffer was added to the ‘input’ sample. 20 µl of 5 M NaCl and 10 µl of RNase A [Roche; 2 mg/ml in 10 mM Tris-HCl (pH7.5) and 15 mM NaCl] were added to each sample and incubated for overnight at 65°C. After incubating for 16 hours, 2 µl of Proteinase K (New England BioLabs), 10 µl of 500 mM EDTA, and 20 µl of 1 M Tris-HCl (pH8.0) were added to each sample and incubated at 45°C for 2 hours. The precipitated DNA was purified using QIAGEN’s PCR purification kit. Real-Time PCR was conducted to quantify precipitated DNA using the Standard Curve method. Power SYBR Green PCR Master Mix (appliedbiosystems) was used as the PCR reaction buffer. The QuantStudio™ 6 Flex System (appliedbiosystems) was used for Real-Time PCR reaction and analyzing the data. Primers used for Real-Time PCR are listed in table S7.

### Fertility assay

Newly eclosed single males (control (*nos-gal4*) or *nos-gal4>UAS-indra*^*TRiP*.*HMJ30228*^) were individually crossed to three *yw* virgin females. After 5 days, each male was transferred to a new vial with three new virgin females. The number of adult flies that eclosed in each vial was scored. *nos-gal4>UAS-indra*^*TRiP*.*HMJ30228*^ females are completely sterile. Therefore, to examine the fertility of *nos>indra*^*TRiP*.*HMJ30228*^ across generations, newly eclosed *nos-gal4>UAS-indra*^*TRiP*.*HMJ30228*^ males for each generation were crossed to *nos-gal4* females to deplete *indra* in germline in the subsequent generation. Then, newly eclosed single males (control (*nos-gal4*) or *nos-gal4>UAS-indra*^*TRiP*.*HMJ30228*^) at each generation were individually crossed to three *yw* virgin females and the number of adult flies that eclosed in each vial was scored.

### Droplet Digital PCR (ddPCR)

20 pairs of testes/sample were dissected from 0-3 day-old control (*nos-gal4*) or *nos-gal4>UAS-indra*^*TRiP*.*HMJ30228*^ males. Genomic DNA isolation was performed as previously described (*9*). Briefly, the testes were homogenized in 200 µl of buffer A (100 mM Tris-HCl pH8.0, 100 mM EDTA pH8.0, 100 mM NaCl, 0.5% SDS), and then an additional 200 µl of buffer A were added to the homogenate. The homogenate was incubated at 65°C for 30 min. Then 800 µl of LiCl/KAc (2.5:1 mixture of 6 M LiCl and 5 M KAc) was added to the homogenate, and the sample was left on ice for 15 min. Subsequently, the sample was centrifuged at 14,000 rpm for 15 min, and 1 ml of supernatant was transferred to a new tube. The supernatant was mixed with 600 µl of isopropanol and centrifuged at 14,000 rpm for 15 min. The pellet (containing genomic DNA) was washed once in 1 ml of 70% ethanol, air dried for 30min and dissolved in 35 µl of TE buffer. The quality and concentration of genomic DNA were measured on a NANODROP ONE (Thermo Scientific).

30 ng of genomic DNA was used per 20 µL ddPCR reaction for control gene reactions (RpL and Upf1), and 0.3 ng of genomic DNA was used per 20 µL ddPCR reaction for 28S rRNA gene reactions. The primers and probes are listed in table S7. ddPCR reactions were carried out according to the manufacturer’s protocol (Bio-Rad). In short, master mixes containing ddPCR Supermix for Probes (No dUTP) (Bio-Rad), genomic DNA, primer/probe mixes, and *Hind*III-HF restriction enzyme (New England Biolabs) for 28S rRNA gene reactions (no restriction enzyme is needed for the control gene reactions) were prepared in 0.2 mL PCR tubes, and incubated at room temperature for 15 min to allow for restriction enzyme digestion. ddPCR droplets were generated from samples using a QX200 Droplet Generator (Bio-Rad), and droplets then underwent complete PCR cycling on a C100 deep-well thermocycler (Bio-Rad). Droplet fluorescence was read using a QX200 Droplet Reader (Bio-Rad). Sample copy number was determined using Quantasoft software (Bio-Rad). rDNA copy number per genome was determined by 28S sample copy number multiplied by 100 (due to the 100x dilution of genomic DNA in the 28S reaction compared to control reaction) divided by control gene copy number multiplied by the expected number of control gene copies per genome (2 for RpL samples; 1 for Upf1 samples). The 28S rRNA gene copy number values normalized by each control were then averaged to determine 28S copy number for each sample.

### Magnification assay

The experimental design to assay rDNA magnification is shown in fig. S4. The *bb*^*z9*^ allele carries an insufficient rDNA copy number on the X chromosome (*10*), which exhibits a ‘*bobbed*’ cuticle phenotype when combined with the *bb*^*158*^ allele (no rDNA on X chromosome) in females. To induce magnification, the *bb*^*z9*^ allele was combined with a Y chromosome lacking rDNA (*bb*^*z9*^*/Ybb*^*-*^)(‘magnifying condition’). These *bb*^*z9*^*/Ybb*^*-*^ males were crossed to *bb*^*158*^*/FM6* females, and the resulting *bb*^*z9*^*/bb*^*158*^ females were examined for the *bobbed* cuticle phenotype. If magnification occurred, a magnified allele (*bb*^*z9-mag*^) combined with *bb*^*158*^ would produce a wild type cuticle, whereas a non-magnified allele combined with *bb*^*158*^ would show a *bobbed* cuticle phenotype. The frequency of wild type cuticle among total female progeny without FM6 (i.e. *bb*^*z9*^ and *bb*^*z9-mag*^ / *bb*^*158*^) was scored as ‘magnification frequency’.

### Statistical analysis

For comparison of sister chromatid segregation patterns in Fig. 1, C and D, Fig. 2F, Fig. 4G, and fig. S1, significance was determined by two-sided Fisher’s exact tests. For comparison of frequencies of *bobbed* animals in Fig. 3B and wild-type cuticle animals in Fig. 3E, significance was determined by two-tailed chi-squared tests using a 2 × 2 contingency table (normal; *bobbed*). Other than these, significance was determined by two-tailed Mann-Whitney tests.

## Supplementary Text

The results shown in Fig. 4 have a few critical implications. Under non-magnifying conditions, USCE appears to be rare, based on the equal amount of IGS FISH signal in anaphase GSCs (Fig. 4, C and E) and based on the fact that both 359-bp repeats and (TAGA)_n_ repeats segregate non-randomly (Fig. 4G). It is interesting to note that without asymmetry in rDNA copy number (under non-magnifying conditions), GSCs still faithfully retain a specific strand (‘red strand’) (Fig. 4, F and G). This suggests that NRSS is not mediated by actual copy number differences, but rather implies that sister chromatids (of rDNA loci) may have additional inherent asymmetries. We speculate that such asymmetries may correlate with the propensity of a specific sister chromatid to gain rDNA copy number, should USCE occur. An attractive candidate for such an asymmetry is the molecular asymmetry during DNA replication that is specific to rDNA loci. It is well established that DNA replication occurs unidirectionally in rDNA loci (*11, 12*) due to the presence of a replication fork block on one side of the replication origin (fig. S5A). Accordingly, one sister chromatid is mostly replicated as the leading strand, whereas the other is mostly replicated as the lagging strand. In yeast, the DNA break that induces rDNA copy number recovery is known to occur on the leading strand (the strand mostly replicated as the lagging strand) at the replication fork block (fig. S5B; (*13*)). If this is universal, the broken end of the leading strand has limited choices as to where to recombine with the sister chromatid to repair the DNA break. The broken end would not recombine with a region that is not yet replicated. The recently replicated region of the lagging strand, where Okazaki fragments have not been processed, may not be a good substrate for sister chromatid recombination, either. The remaining possible region would be the sister chromatid that was replicated as the leading strand (fig. S5B). If this happens, the strand mostly replicated as the lagging strand is likely to gain the copy number. Thus, we speculate that the mechanism that mediates NRSS may have the ability to distinguish leading vs. lagging strands and specifically connects the lagging strand to the GSC side.

Although the mechanism that ensures the retention of a specific strand to the GSC remains elusive, the CO-FISH results shown in Fig. 4, F and G provide a critical hint. Under magnifying conditions, where USCE occurs, it is the (TAGA)_n_ repeats whose segregation pattern is randomized. This suggests that the 359-bp side of the rDNA is responsible for the retention in GSCs. This side of the chromosome contains the centromere, whose asymmetry has been suggested to mediate non-random segregation of chromosomes (*14, 15*). However, we have no evidence thus far to suggest that the centromere is responsible for NRSS of the X and Y chromosomes. Additionally, the loss of rDNA with retention of most of 359-bp and the entire centromere was sufficient to compromise NRSS (Fig. 1C). Therefore, it is highly unlikely that 359-bp or the centromere contains sufficient information to mediate NRSS. Future investigation is required to address these key molecular mechanisms.

**Fig. S1.**
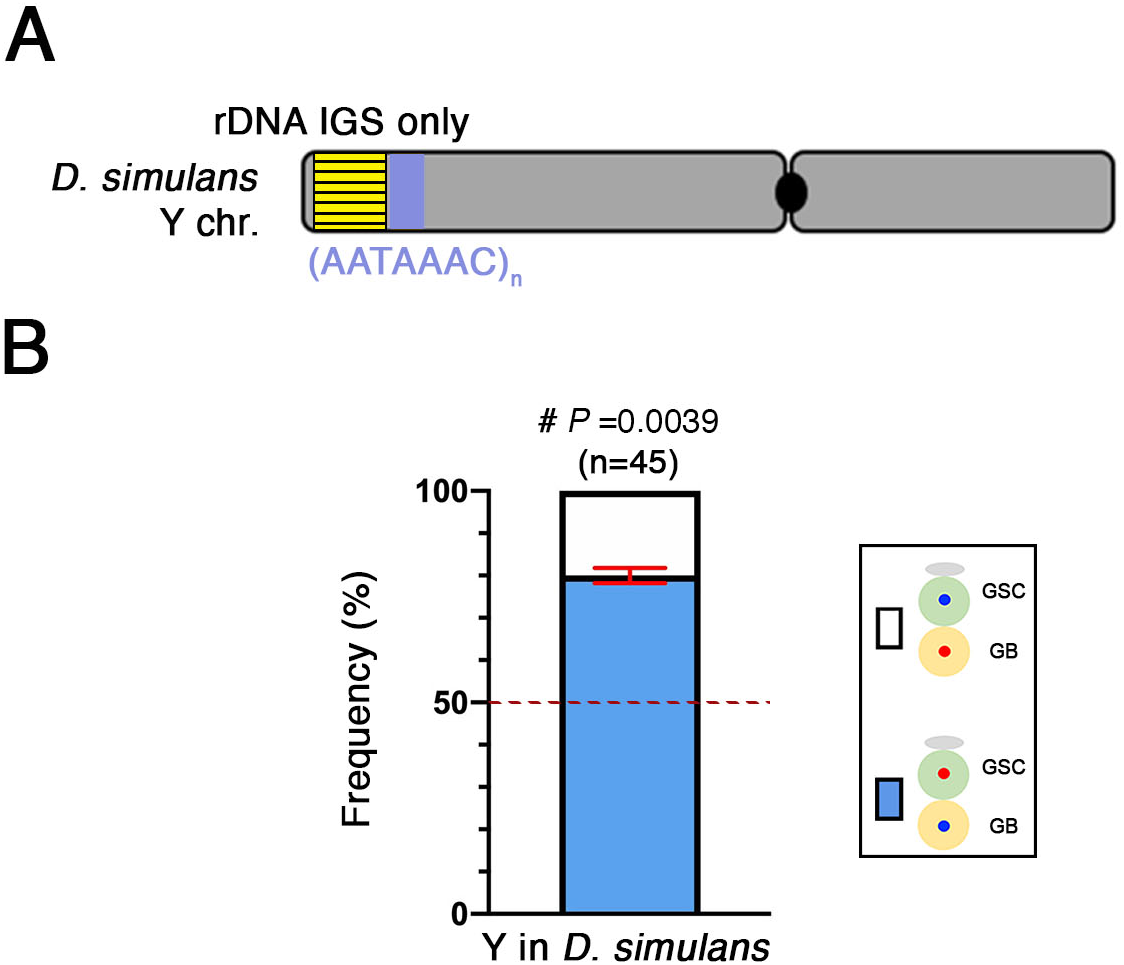
The *D. simulans* Y chromosome segregates sister chromatids non-randomly. (A) Schematic of the *D. simulans* Y chromosome (B) Summary of the sister chromatid segregation pattern assessed by CO-FISH in the *D. simulans* control strain (*w*^*501*^) (see table S1 for detailed data). Data shown as mean ± s.d. from three independent experiments. n, number of GSC-GB pairs scored. #, *P*-value of Fisher’s exact test by comparing to hypothetical random sister chromatid segregation is shown.

**Fig. S2.**
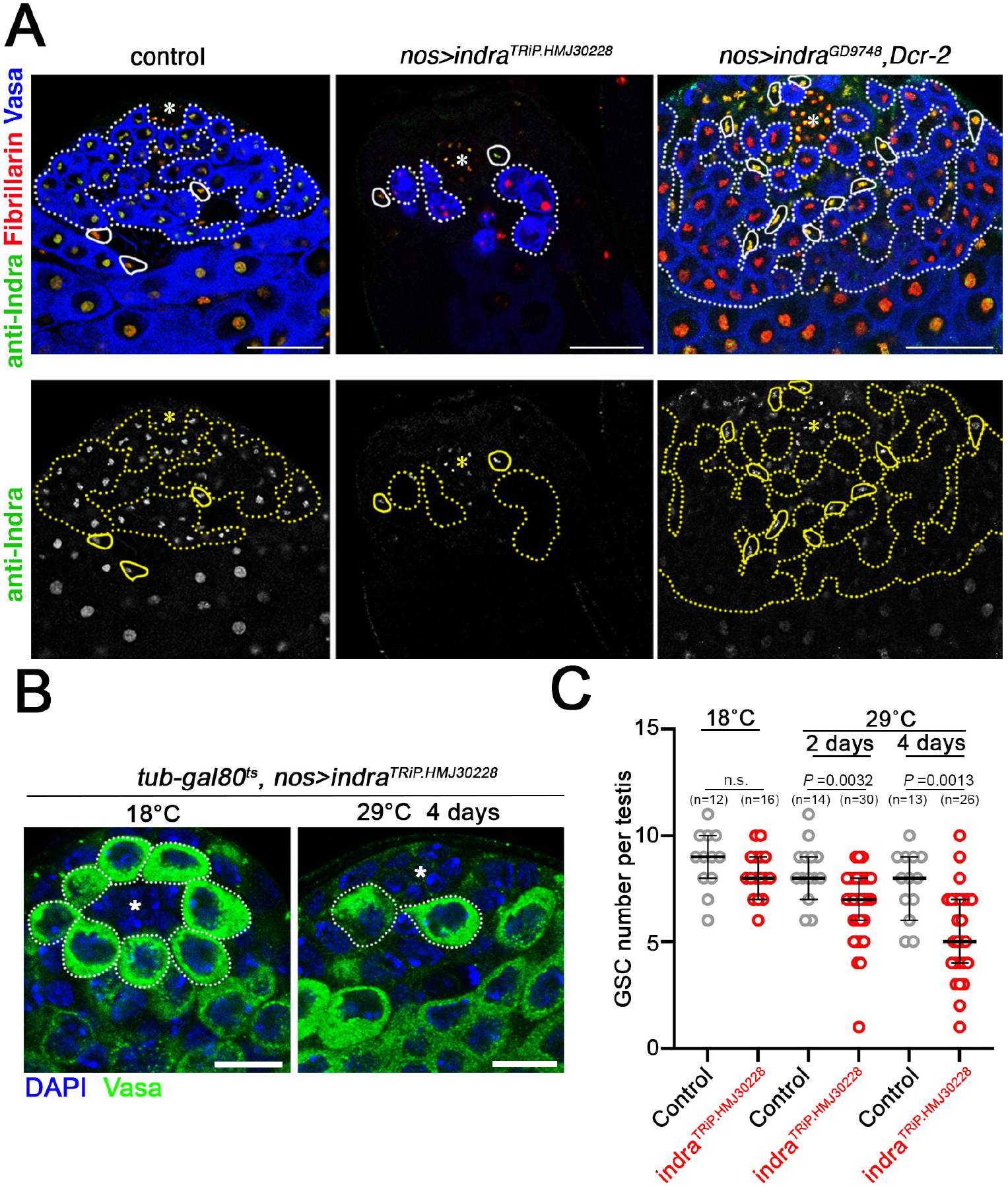
Validation of *indra*^*RNAi*^ efficiency and antibody specificity. (A) Examples of testis apical tips after *indra* knockdown by indicated *indra*^*RNAi*^ lines. Anti-Indra antibody staining was lost from germ cells upon *indra* knockdown by *nos-gal4>indra*^*TRiP*.*HMJ30228*^ or *nos-gal4>indra*^*GD9748*^ *dcr-2*. (UAS-dcr-2 was added to enhance the efficiency of *indra*^*GD9748*^). This experiment also demonstrates the specificity of the anti-Indra antibody. The hub is indicated by an asterisk. Germ cells are indicated by dotted lines and somatic cells are indicated by solid lines. Bar: 25 µm. (B) Examples of testis apical tips before and after induction of *nos-gal4>indra*^*TRiP*.*HMJ30228*^. GSCs are indicated by dotted lines and the hub is indicated by an asterisk. Bar: 10 µm. (C) GSC number after induction of *indra*^*TRiP*.*HMJ30228*^. n, number of testes scored. *P*-values: two-tailed Mann-Whitney test.

**Fig. S3.**
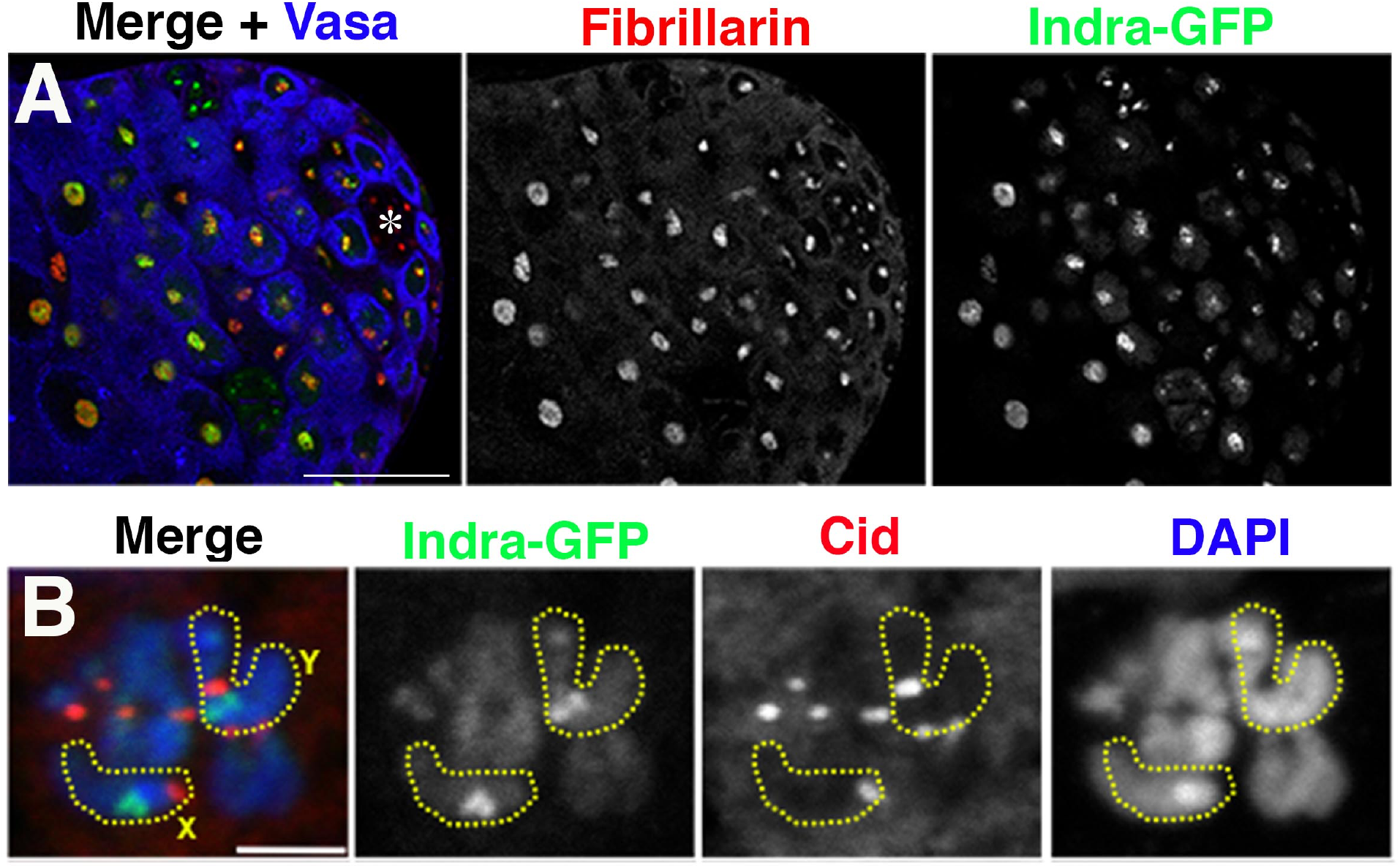
Localization of Indra-GFP to the nucleolus and rDNA loci. (A) Localization of Indra-GFP at the apical tip of the testis. Indra localizes to nucleolus visualized by Fibrillarin. The hub is indicated by an asterisk. Bar: 25 µm. (B) Localization of Indra-GFP on a metaphase chromosome spread. The X and Y chromosomes are indicated by dotted lines. Cid: centromere. Bar: 5 µm.

**Fig. S4.**
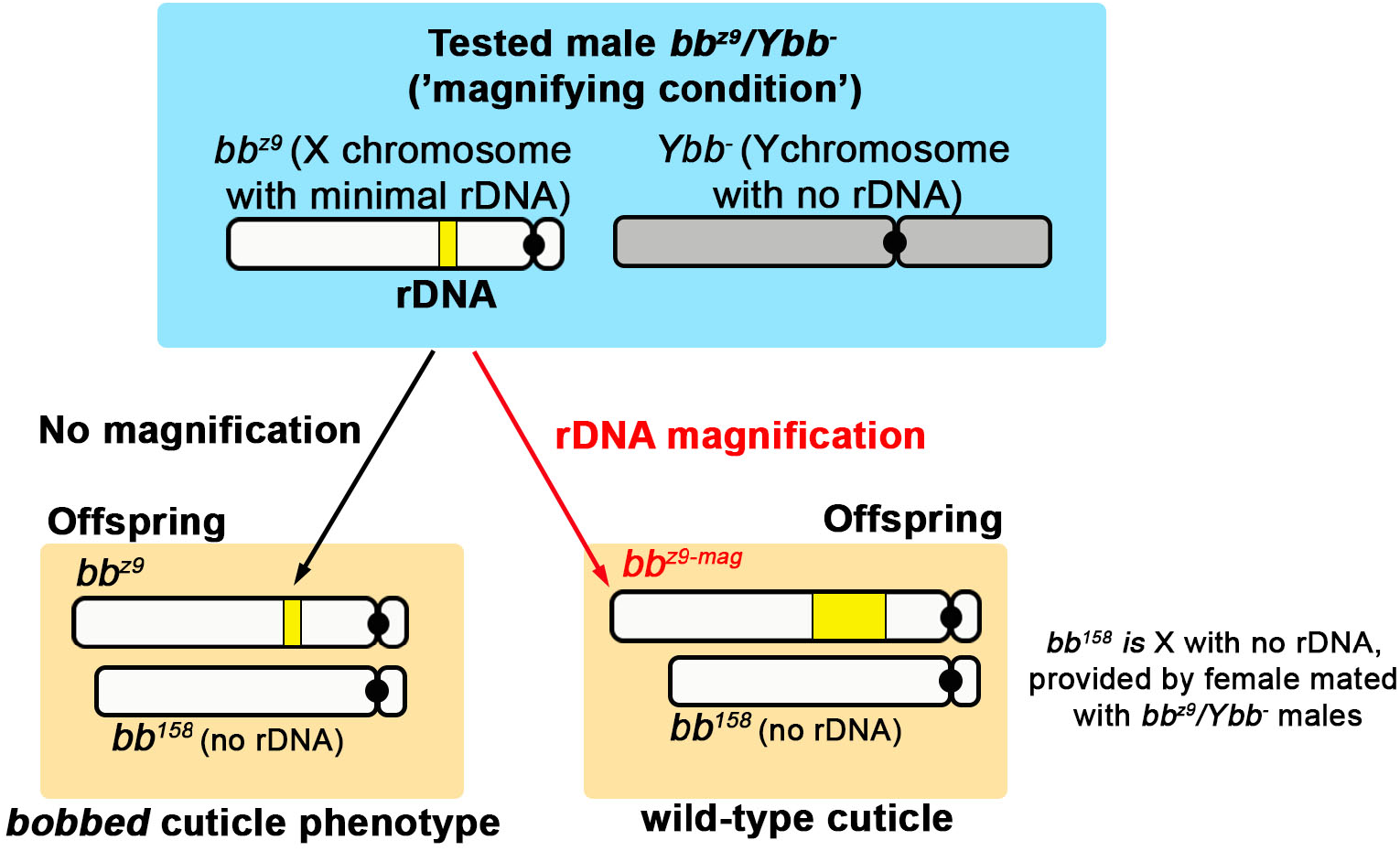
Diagram of phenotypic assessment to detect rDNA magnification of the bb^z9^ allele. The *bb*^*z9*^ allele carries an insufficient rDNA copy number on the X chromosome, which causes flies to exhibit a ‘*bobbed*’ cuticle phenotype when combined with the *bb*^*158*^ allele (no rDNA on X chromosome) in females (Fig. 3B). To induce magnification, the *bb*^*z9*^ allele was combined with a Y chromosome without rDNA (*bb*^*z9*^*/Ybb*^*-*^)(‘magnifying condition’). To assess whether the *bb*^*z9*^ allele magnified, these *bb*^*z9*^*/Ybb*^*-*^ males were crossed to *bb*^*158*^*/FM6* female, and cuticle phenotype of the resulting *bb*^*z9*^*/bb*^*158*^ females was examined. If magnification occurred, the magnified allele (*bb*^*z9-mag*^) combined with *bb*^*158*^ would have a wild type cuticle, whereas the non-magnified allele combined with *bb*^*158*^ would have the *bobbed* phenotype. The frequency of wild type cuticle among total female progeny without FM6 (i.e. *bb*^*z9*^ and *bb*^*z9-mag*^/*bb*^*158*^) was scored as ‘magnification frequency’.

**Fig. S5.**
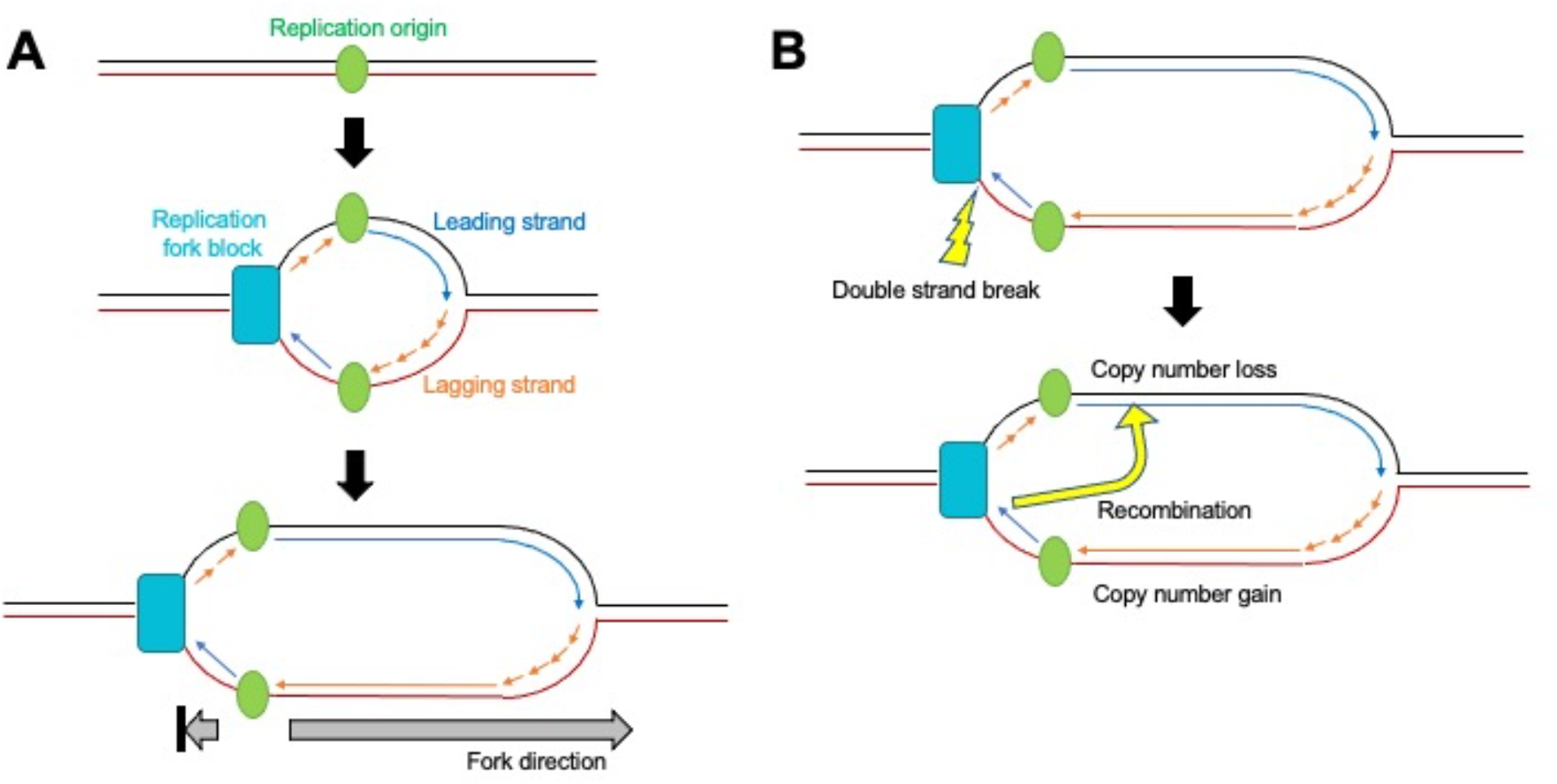
Diagram of DNA replication at rDNA loci. (A) Replication fork block on one side of the replication origin leads to mostly unidirectional DNA replication at the rDNA loci. This causes one sister chromatid to be synthesized primarily as leading strand and the other as lagging strand. (B) In yeast, double strand DNA breaks primarily occur on the leading strand when fork progression is prevented at the replication fork block (top). An appropriate donor for DNA repair may be found in the region of the sister chromatid replicated as leading strand. If such recombination happens, the sister chromatid mostly replicated as lagging strand (bottom strand) will gain copy number.

**Table S1.**
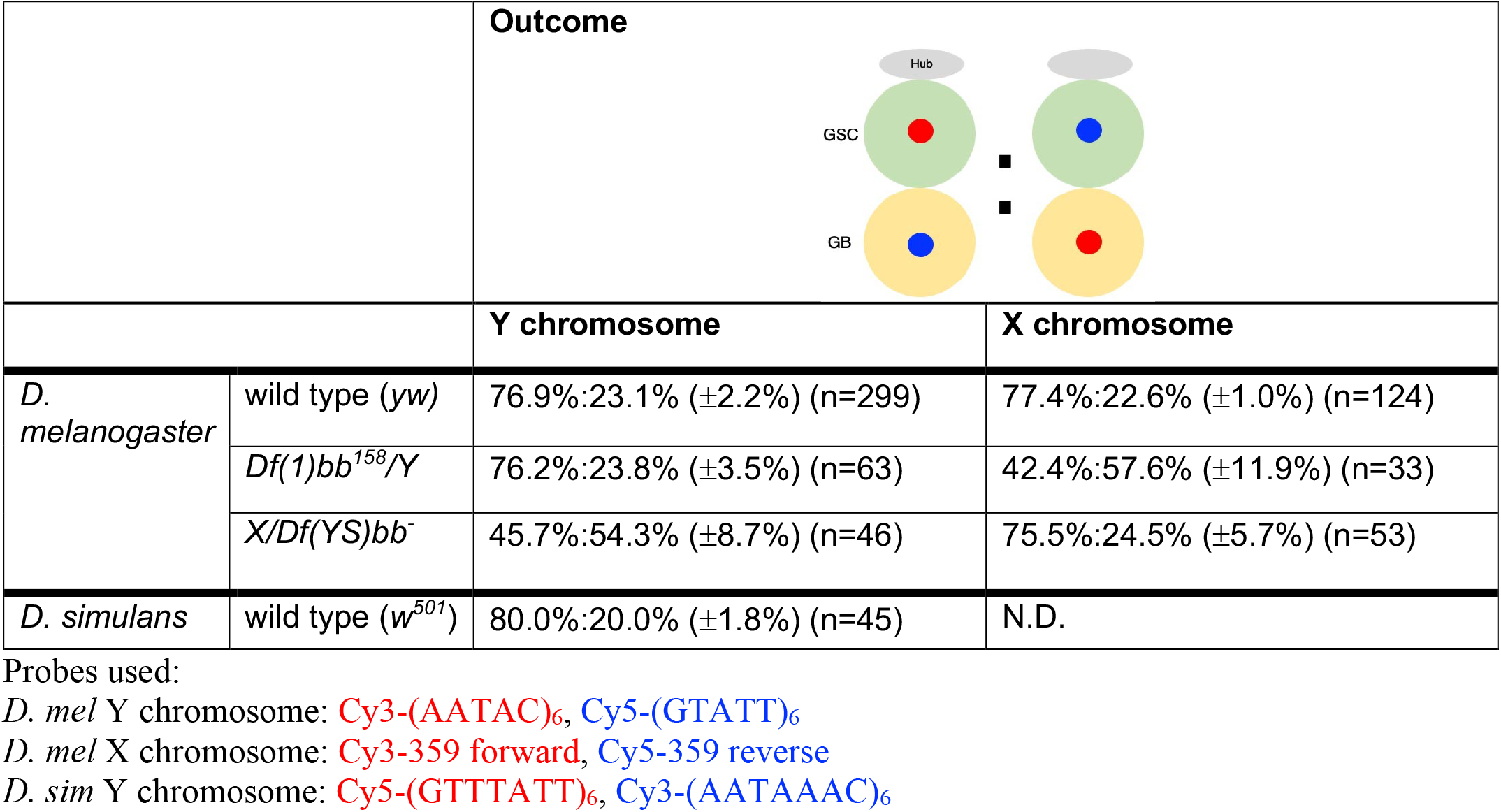
CO-FISH results in *D. melanogaster* rDNA deficient stocks and *D. simulans*.

|  |  | **Outcome** |  |
| --- | --- | --- | --- |
|  |  | **Y chromosome** | **X chromosome** |
| *D. melanogaster* | wild type (*yw*) | 76.9%:23.1% ($\pm 2.2\%$) (n=299) | 77.4%:22.6% ($\pm 1.0\%$) (n=124) |
| | *Df(1)bb<sup>158</sup>/Y* | 76.2%:23.8% ($\pm 3.5\%$) (n=63) | 42.4%:57.6% ($\pm 11.9\%$) (n=33) |
| | *X/Df(YS)bb<sup>-</sup>* | 45.7%:54.3% ($\pm 8.7\%$) (n=46) | 75.5%:24.5% ($\pm 5.7\%$) (n=53) |
| *D. simulans* | wild type (*w<sup>501</sup>*) | 80.0%:20.0% ($\pm 1.8\%$) (n=45) | N.D. |

Probes used: *D. mel* Y chromosome: Cy3-(AATAC)<sub>6</sub>, Cy5-(GTATT)<sub>6</sub> *D. mel* X chromosome: Cy3-359 forward, Cy5-359 reverse *D. sim* Y chromosome: Cy5-(GTTTATT)<sub>6</sub>, Cy3-(AATAAAC)<sub>6</sub>

**Table S2.**
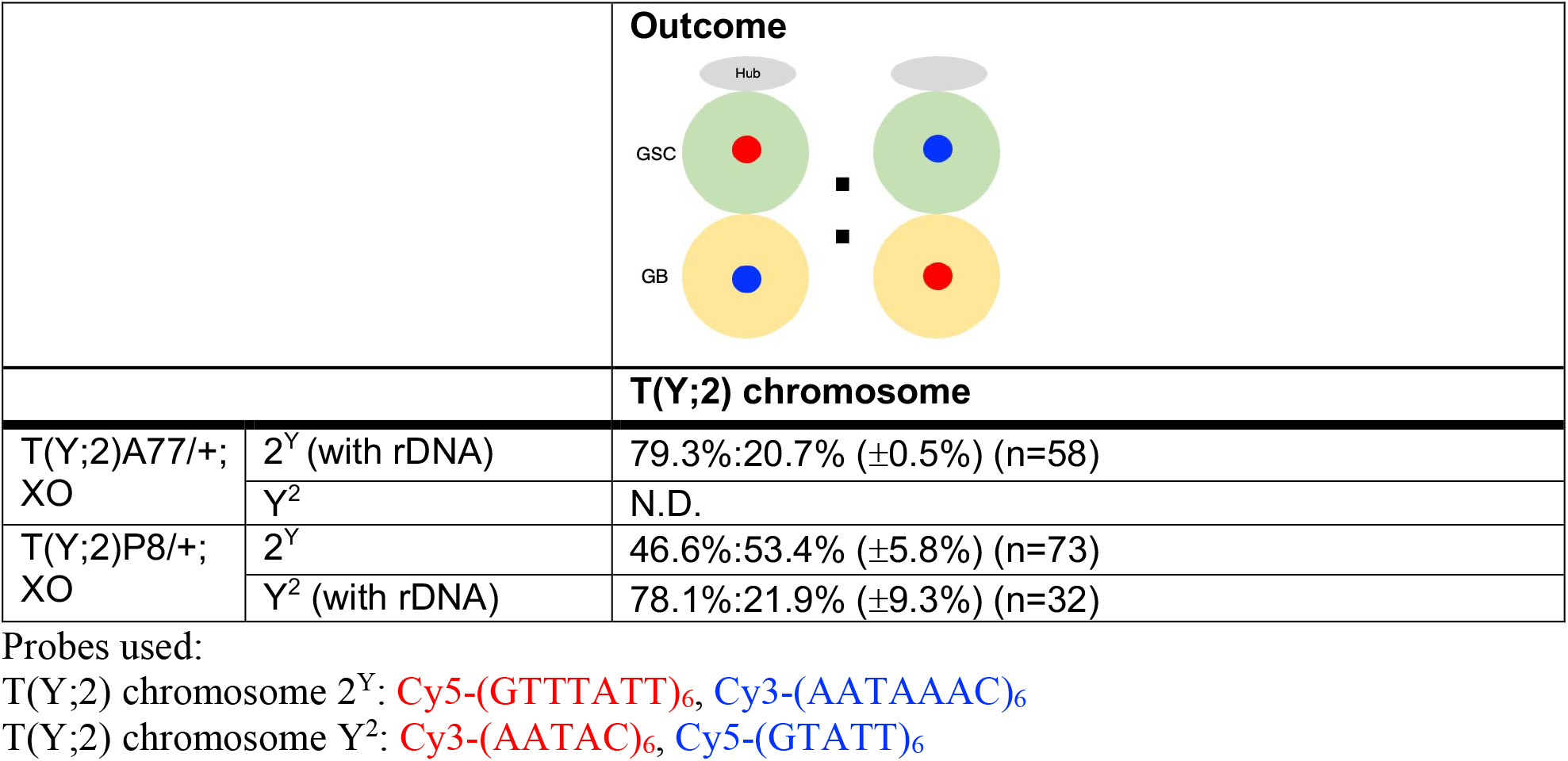
CO-FISH results of Y-2 translocation chromosomes.

|  |  | **Outcome** |
| --- | --- | --- |
|  |  | **T(Y;2) chromosome** |
| T(Y;2)A77/+;<br>XO | 2<sup>Y</sup> (with rDNA) | 79.3%:20.7% ($\pm 0.5\%$) (n=58) |
|  | Y<sup>2</sup> | N.D. |
| T(Y;2)P8/+;<br>XO | 2<sup>Y</sup> | 46.6%:53.4% ($\pm 5.8\%$) (n=73) |
| | Y<sup>2</sup> (with rDNA) | 78.1%:21.9% ($\pm 9.3\%$) (n=32) |

Probes used: T(Y;2) chromosome 2<sup>Y</sup>: Cy5-(GTTTATT)<sub>6</sub>, Cy3-(AATAAAC)<sub>6</sub> T(Y;2) chromosome Y<sup>2</sup>: Cy3-(AATAC)<sub>6</sub>, Cy5-(GTATT)<sub>6</sub>

**Table S3.**
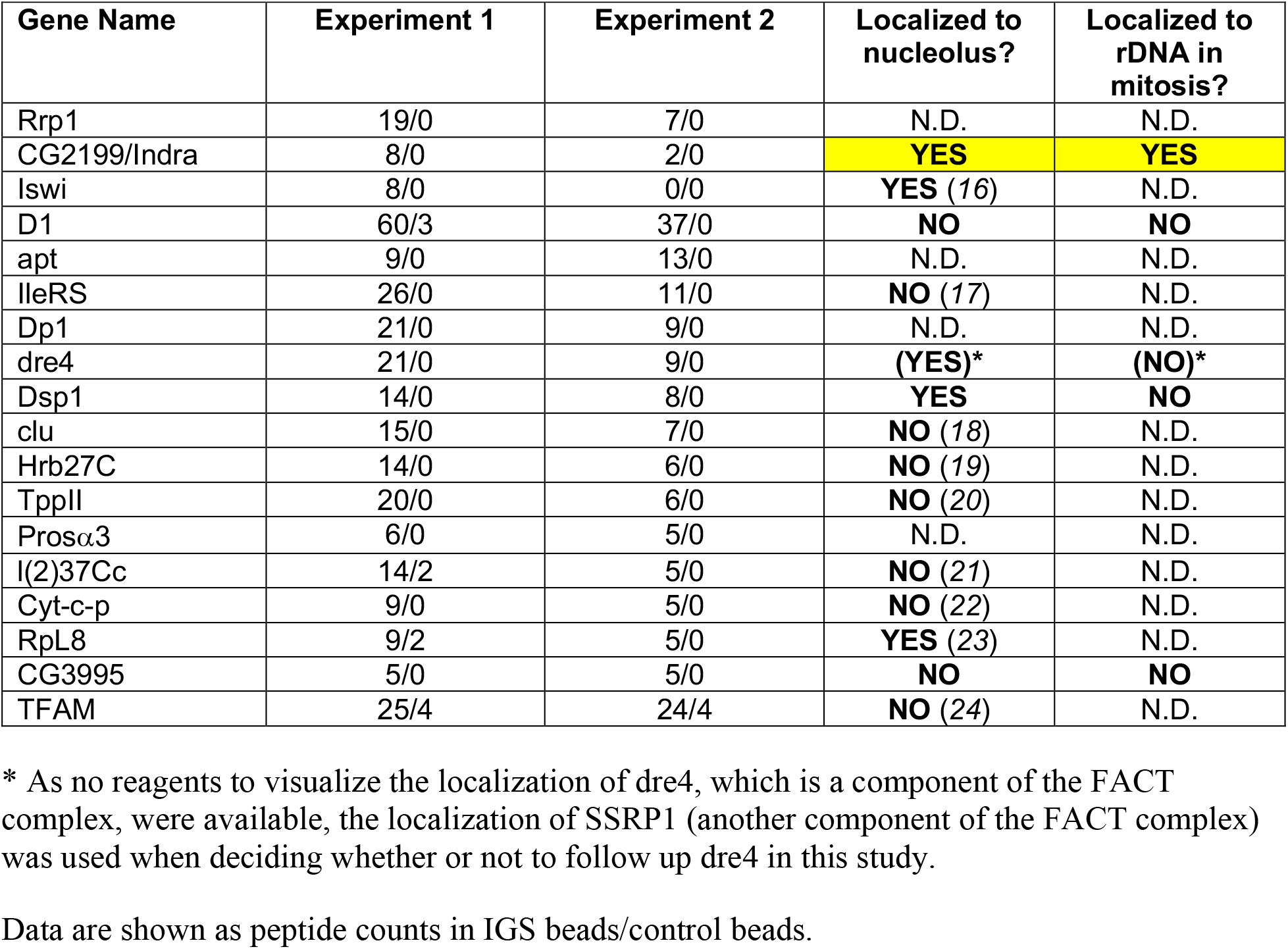
List of proteins that were enriched in IGS-beads pull-down.

**Table S4.** CO-FISH results upon knockdown of *indra*.

|  | Outcome |  |
| --- | --- | --- |
|  | Y chromosome | X chromosome |
| *indra*<sup>TRiP.HMJ30228</sup> control* | 75.0%:25.0% ($\pm 2.6\%$) (n=40) | 80.0%:20.0% ($\pm 1.7\%$) (n=30) |
| *tub-gal80<sup>ts</sup>, nos-gal4ΔVP16>UAS-indra<sup>HMJ30228</sup>* | 40.0%:60.0% ($\pm 0.0\%$) (n=30) | 43.6%:56.4% ($\pm 9.6\%$) (n=39) |
| *indra*<sup>GD9748</sup> control<br>(*nos-gal4>UAS-Dcr-2*) | 76.9%:23.1% ($\pm 7.4\%$) (n=26) | 76.9%:23.1% ($\pm 7.9\%$) (n=39) |
| *nos-gal4>UAS-indra*<sup>GD9748</sup>,<br>*UAS-Dcr-2* | 42.9%:57.1% ($\pm 2.1\%$) (n=28) | 43.2%:56.8% ($\pm 7.5\%$) (n=44) |

Probes used:
Y chromosome: **Cy3-(AATAC)<sub>6</sub>**, **Cy5-(GTATT)<sub>6</sub>**
X chromosome: **Cy3-359 forward**, **Cy5-359 reverse**
\*: Cross siblings of *tub-gal80<sup>ts</sup>, nos-gal4 $\Delta$ VP16>UAS-indra<sup>TRiP.HMJ30228</sup>* that do not express *indra*<sup>HMJ30228</sup> (either *nos-gal4 $\Delta$ VP16* only or *UAS-indra<sup>TRiP.HMJ30228</sup>* only) were used as control

**Table S5.**
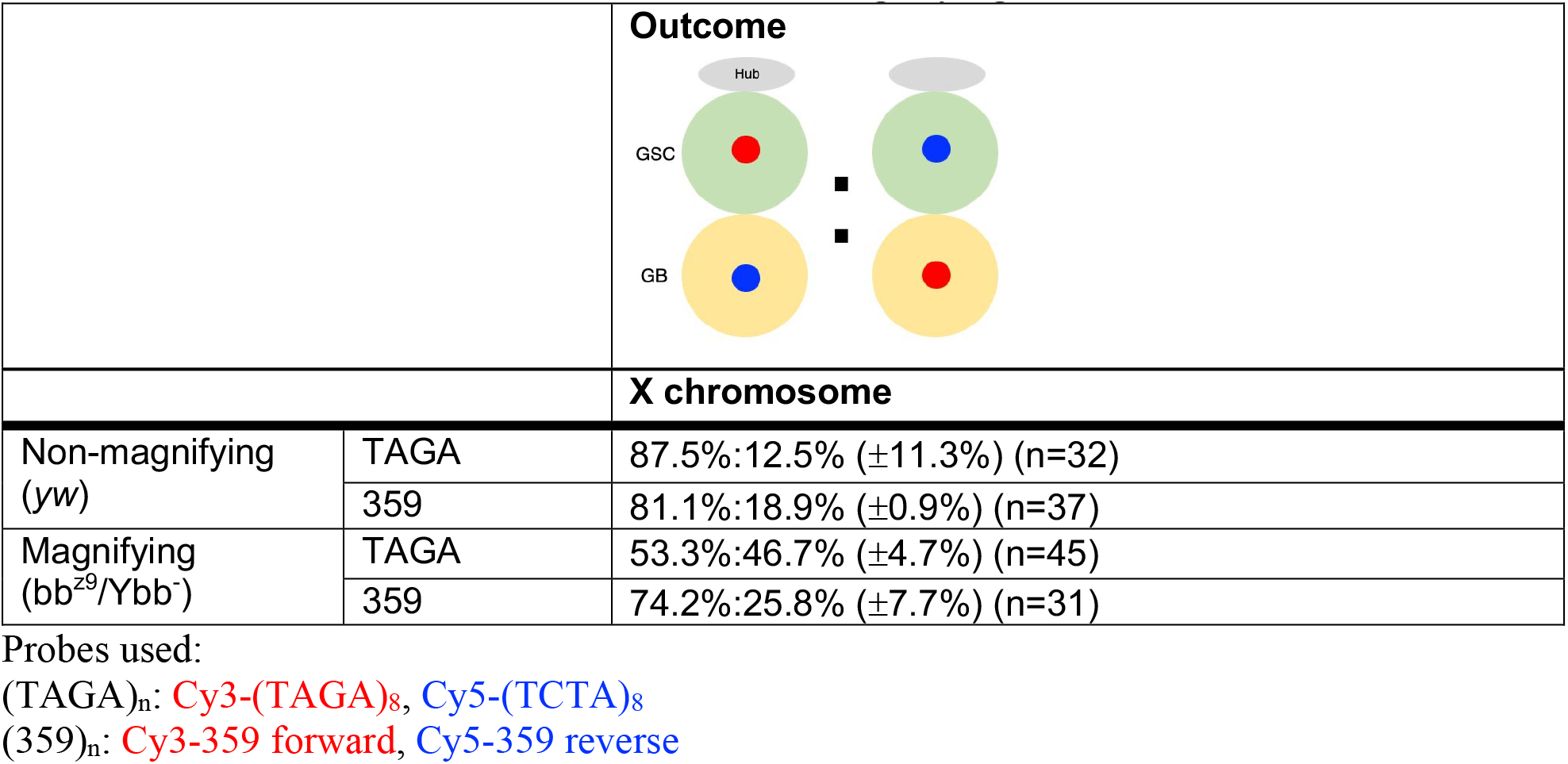
CO-FISH results of X chromosome in magnifying condition.

**Table S6.**
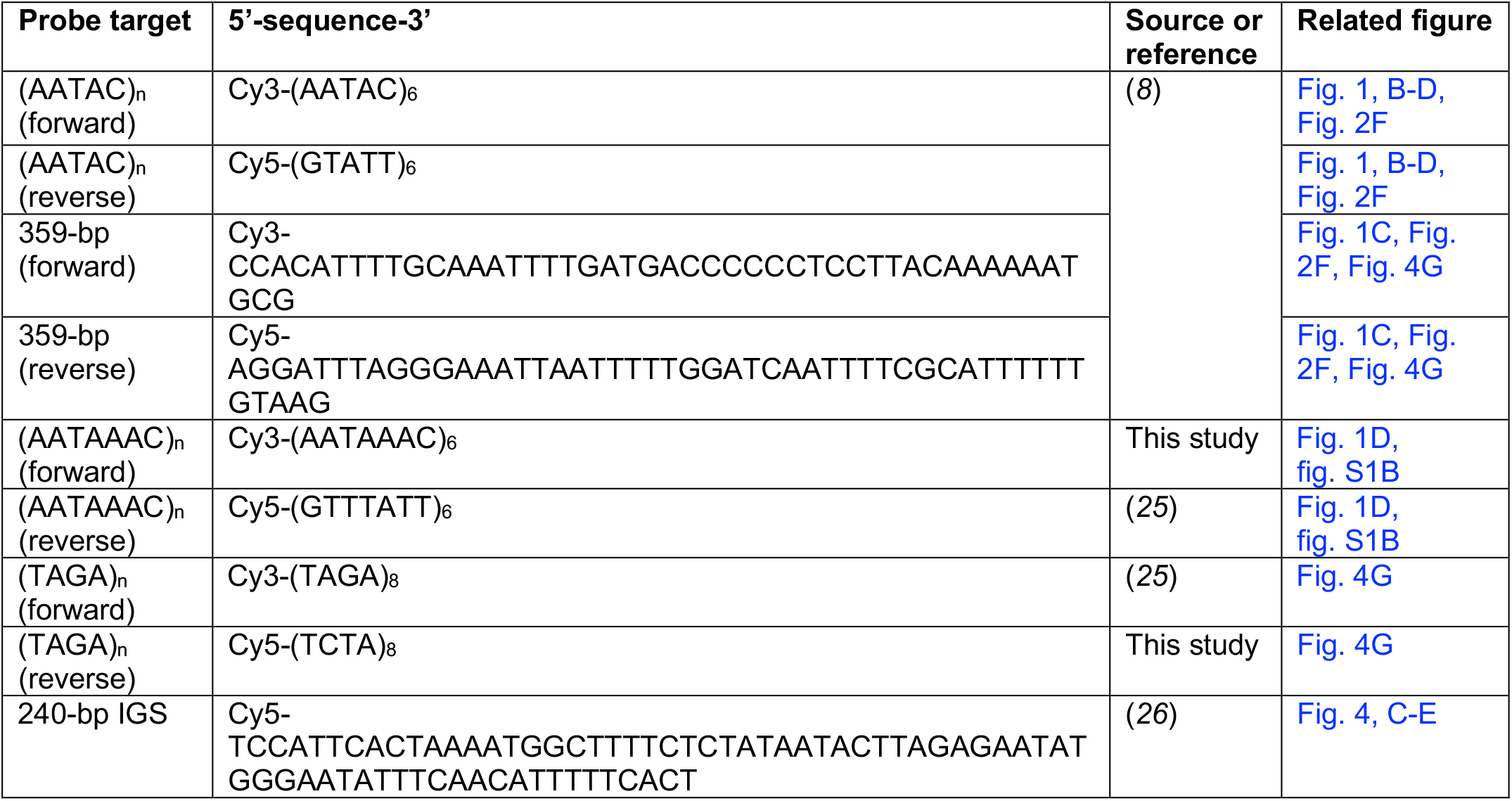
Probe sequences for CO-FISH and DNA FISH.

**Table S7:**
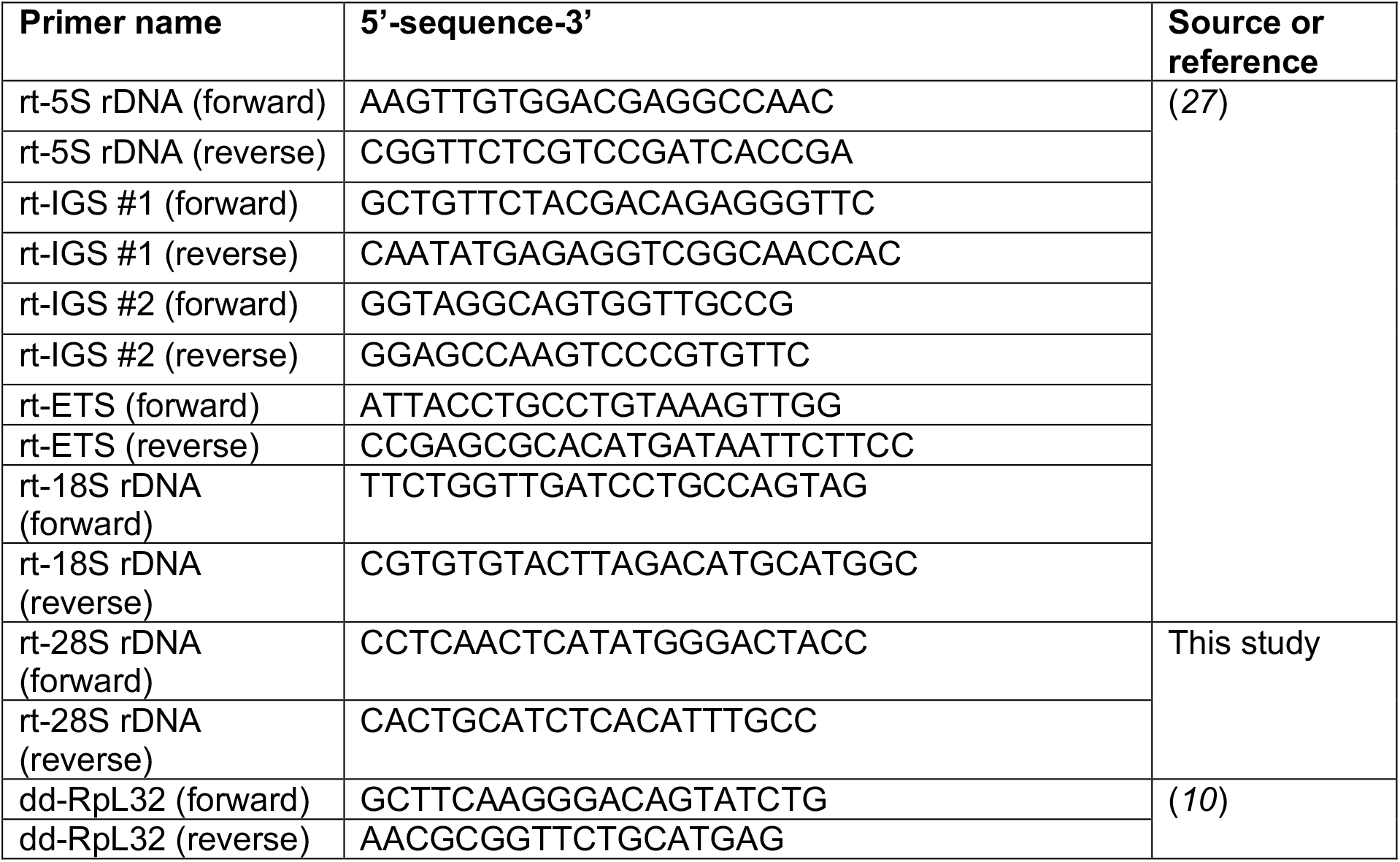
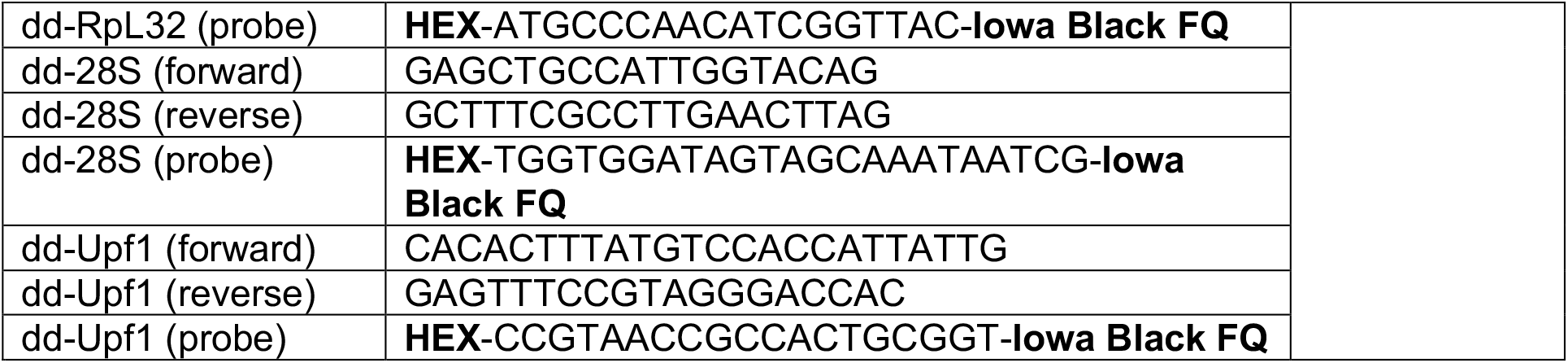
Primer and probe sequences for Real-Time PCR and Droplet-Digital PC.

## Notes

### Competing Interest Statement

The authors have declared no competing interest.

## References

1. P. M. Lansdorp, Immortal strands? Give me a break. Cell 129, 1244–1247 (2007).

2. T. A. Rando, The immortal strand hypothesis: segregation and reconstruction. Cell 129, 1239–1243 (2007).

3. S. Tajbakhsh, C. Gonzalez, Biased segregation of DNA and centrosomes: moving together or drifting apart? Nat Rev Mol Cell Biol 10, 804–810 (2009).

4. S. Yadlapalli, Y. M. Yamashita, Chromosome-specific nonrandom sister chromatid segregation during stem-cell division. Nature 498, 251–254 (2013).

5. S. Tajbakhsh, Stem cell identity and template DNA strand segregation. Curr Opin Cell Biol 20, 716–722 (2008).

6. R. S. Hawley, C. H. Marcus, Recombinational controls of rDNA redundancy in Drosophila. Annu Rev Genet 23, 87–120 (1989).

7. A. R. Lohe, P. A. Roberts, An unusual Y chromosome of Drosophila simulans carrying amplified rDNA spacer without rRNA genes. Genetics 125, 399–406 (1990).

8. F. M. Ritossa, K. C. Atwood, S. Spiegelman, A molecular explanation of the bobbed mutants of Drosophila as partial deficiencies of “ribosomal” DNA. Genetics 54, 819–834 (1966).

9. K. D. Tartof, Unequal mitotic sister chromatin exchange as the mechanism of ribosomal RNA gene magnification. Proc Natl Acad Sci U S A 71, 1272–1276 (1974).

10. T. Kobayashi, Ribosomal RNA gene repeats, their stability and cellular senescence. Proc Jpn Acad Ser B Phys Biol Sci 90, 119–129 (2014).

11. K. L. Lu, J. O. Nelson, G. J. Watase, N. Warsinger-Pepe, Y. M. Yamashita, Transgenerational dynamics of rDNA copy number in Drosophila male germline stem cells. Elife 7, (2018).

## References

1. M. Van Doren, A. L. Williamson, R. Lehmann, Regulation of zygotic gene expression in Drosophila primordial germ cells. Curr Biol 8, 243–246 (1998).

2. M. P. Zeidler, N. Perrimon, D. I. Strutt, Polarity determination in the Drosophila eye: a novel role for unpaired and JAK/STAT signaling. Genes Dev 13, 1342–1353 (1999).

3. S. E. McGuire, P. T. Le, A. J. Osborn, K. Matsumoto, R. L. Davis, Spatiotemporal rescue of memory dysfunction in Drosophila. Science 302, 1765–1768 (2003).

4. M. Inaba, M. Buszczak, Y. M. Yamashita, Nanotubes mediate niche-stem-cell signalling in the Drosophila testis. Nature 523, 329–332 (2015).

5. M. Zaccai, H. D. Lipshitz, Differential distributions of two adducin-like protein isoforms in the Drosophila ovary and early embryo. Zygote 4, 159–166 (1996).

6. B. Riggleman, P. Schedl, E. Wieschaus, Spatial expression of the Drosophila segment polarity gene armadillo is posttranscriptionally regulated by wingless. Cell 63, 549–560 (1990).

7. M. Jagannathan, R. Cummings, Y. M. Yamashita, A conserved function for pericentromeric satellite DNA. Elife 7, (2018).

8. S. Yadlapalli, Y. M. Yamashita, Chromosome-specific nonrandom sister chromatid segregation during stem-cell division. Nature 498, 251–254 (2013).

9. A. M. Huang, E. J. Rehm, G. M. Rubin, Quick preparation of genomic DNA from Drosophila. Cold Spring Harb Protoc 2009, pdb prot5198 (2009).

10. J. O. Nelson, A. Slicko, Y. M. Yamashita, The retrotransposon R2 maintains Drosophila ribosomal DNA repeats. bioRxiv, 2021.2007.2012.451825 (2021).

11. Y. Akamatsu, T. Kobayashi, The Human RNA Polymerase I Transcription Terminator Complex Acts as a Replication Fork Barrier That Coordinates the Progress of Replication with rRNA Transcription Activity. Mol Cell Biol 35, 1871–1881 (2015).

12. T. Kobayashi, Ribosomal RNA gene repeats, their stability and cellular senescence. Proc Jpn Acad Ser B Phys Biol Sci 90, 119–129 (2014).

13. M. D. Burkhalter, J. M. Sogo, rDNA enhancer affects replication initiation and mitotic recombination: Fob1 mediates nucleolytic processing independently of replication. Mol Cell 15, 409–421 (2004).

14. T. Akera, E. Trimm, M. A. Lampson, Molecular Strategies of Meiotic Cheating by Selfish Centromeres. Cell 178, 1132–1144 e1110 (2019).

15. R. Ranjan, J. Snedeker, X. Chen, Asymmetric Centromeres Differentially Coordinate with Mitotic Machinery to Ensure Biased Sister Chromatid Segregation in Germline Stem Cells. Cell Stem Cell 25, 666–681 e665 (2019).

16. A. V. Emelyanov et al., Identification and characterization of ToRC, a novel ISWI-containing ATP-dependent chromatin assembly complex. Genes Dev 26, 603–614 (2012).

17. J. Lu, S. J. Marygold, W. H. Gharib, B. Suter, The aminoacyl-tRNA synthetases of Drosophila melanogaster. Fly (Austin) 9, 53–61 (2015).

18. R. T. Cox, A. C. Spradling, Clueless, a conserved Drosophila gene required for mitochondrial subcellular localization, interacts genetically with parkin. Dis Model Mech 2, 490–499 (2009).

19. M. J. Matunis, E. L. Matunis, G. Dreyfuss, Isolation of hnRNP complexes from Drosophila melanogaster. J Cell Biol 116, 245–255 (1992).

20. S. C. Renn, B. Tomkinson, P. H. Taghert, Characterization and cloning of tripeptidyl peptidase II from the fruit fly, Drosophila melanogaster. J Biol Chem 273, 19173–19182 (1998).

21. S. J. Lee, R. Feldman, P.H. O’Farrell, An RNA interference screen identifies a novel regulator of target of rapamycin that mediates hypoxia suppression of translation in Drosophila S2 cells. Mol Biol Cell 19, 4051–4061 (2008).

22. L. Dorstyn, K. Mills, Y. Lazebnik, S. Kumar, The two cytochrome c species, DC3 and DC4, are not required for caspase activation and apoptosis in Drosophila cells. J Cell Biol 167, 405–410 (2004).

23. K. N. Rugjee et al., Fluorescent protein tagging confirms the presence of ribosomal proteins at Drosophila polytene chromosomes. PeerJ 1, e15 (2013).

24. K. Takata et al., Drosophila mitochondrial transcription factor A: characterization of its cDNA and expression pattern during development. Biochem Biophys Res Commun 287, 474–483 (2001).

25. M. Jagannathan, N. Warsinger-Pepe, G. J. Watase, Y. M. Yamashita, Comparative Analysis of Satellite DNA in the Drosophila melanogaster Species Complex. G3 (Bethesda) 7, 693–704 (2017).

26. K. L. Lu, J. O. Nelson, G. J. Watase, N. Warsinger-Pepe, Y. M. Yamashita, Transgenerational dynamics of rDNA copy number in Drosophila male germline stem cells. Elife 7, (2018).

27. Q. Zhang, N. A. Shalaby, M. Buszczak, Changes in rRNA transcription influence proliferation and cell fate within a stem cell lineage. Science 343, 298–301 (2014).

